# Membrane-permeable 5-fluorodeoxyuridine triphosphate derivatives inhibit the proliferation of *Plasmodium falciparum*

**DOI:** 10.1101/2025.02.25.640175

**Authors:** Vella Nikolova, Karen Linnemannstöns, Marie-Elise Bendel, Marta Machado, Benedikt Ganter, Patricia Budimir, Michelle Vogts, Markus Ganter, Chris Meier, Matthias Dobbelstein

**Affiliations:** Department of Molecular Oncology, Göttingen Center of Molecular Biosciences (GZMB), University Medical Center Göttingen, Justus-von-Liebig-Weg 11, 37077 Göttingen, Germany; Max Planck Institute for Multidisciplinary Sciences, Am Fassberg 11, 37077 Göttingen, Germany; Center for Infectious Diseases – Parasitology, Medical Faculty, Heidelberg University, Im Neuenheimer Feld 324, 69120 Heidelberg, Germany; Organic Chemistry, Department of Chemistry, Faculty of Mathematics, Informatics and Natural Sciences, University of Hamburg, Martin-Luther-King-Platz 6, 20146, Hamburg, Germany; Centre for Structural Systems Biology (CSSB), Hamburg, DESY Campus, Notkestrasse 85, D-22607 Hamburg, Germany

**Keywords:** Malaria, *Plasmodium falciparum*, nucleoside analogues, nucleotides, fluorouridine, 5-fluorodeoxyuridine

## Abstract

Malaria tropica remains a major global health challenge, requiring new therapeutic strategies against *Plasmodium falciparum*. While nucleoside analogues are effective against viruses and cancer, their use against *P. falciparum* is limited by the parasite’s lack of nucleoside kinases. To overcome this, we tested cell-permeable derivatives of 5-fluorodeoxyuridine triphosphate (cpFdUTP) for anti-parasitic activity in infected human red blood cells. cpFdUTP rapidly and potently inhibited P. falciparum proliferation, arresting development at the trophozoite-to-schizont transition by stalling DNA replication, as revealed by a *P. falciparum* nuclear cycle sensor line. Although cpFdUTP also impaired human cell growth, supplementation with thymidine or cell-permeable deoxythymidine triphosphate (cpdTTP) selectively rescued human cells while maintaining P. falciparum inhibition. This identifies a potential therapeutic window for cpFdUTP in combination with thymidine, outlining a novel approach for malaria treatment.

**Graphical Abstract:** 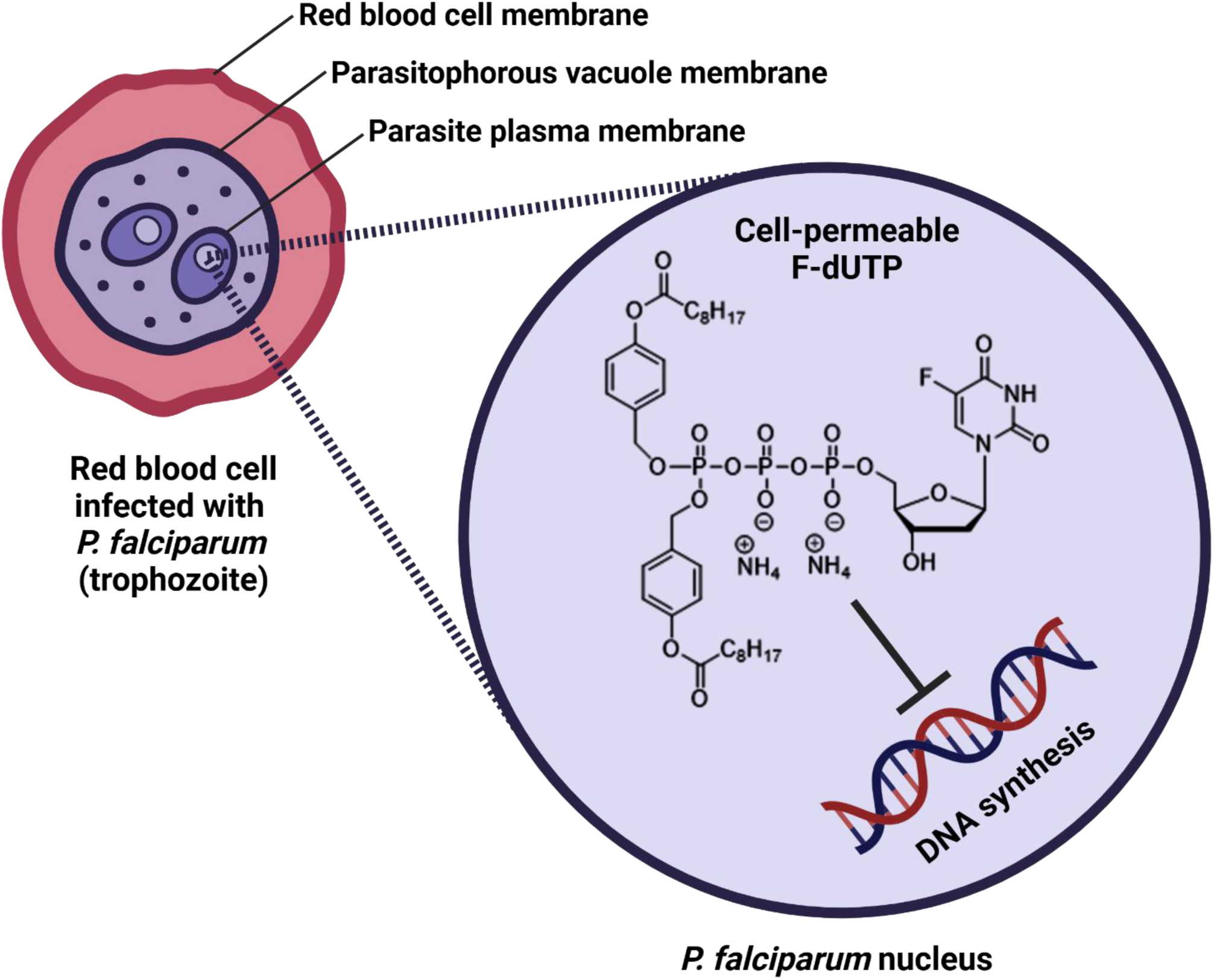

## INTRODUCTION

Malaria remains one of the most devastating infectious diseases globally, causing hundreds of thousands of deaths annually, predominantly among children in sub-Saharan Africa (Phillips et al., 2017). The protozoan parasite *Plasmodium falciparum* (*P. falciparum*) is responsible for the most severe form of malaria in humans. Proliferation of *P. falciparum* inside red blood cells (RBCs) is responsible for all clinical symptoms of the disease (Cowman et al., 2016), and *P. falciparum* has developed resistance to many frontline antimalarial drugs, emphasizing the urgent need for novel therapeutic strategies (Okombo and Fidock, 2024). Incomplete protection by currently available vaccines further underscores this necessity (Duffy et al., 2024). Moreover, current treatment regimens rely heavily on artemisinin-based combination therapies (ACTs), but reduced susceptibility to artemisinin and resistance to its partner drugs is rapidly emerging, posing a significant threat to malaria control and eradication efforts (Rosenthal et al., 2024).

Nucleoside analogues have demonstrated remarkable therapeutic utility in oncology (Galmarini et al., 2002; Tsesmetzis et al., 2018) and antiviral therapies (Seley-Radtke and Yates, 2018; Yates and Seley-Radtke, 2019) by targeting nucleotide biosynthesis and disrupting the assembly of nucleic acids (Jordheim et al., 2013). These compounds act as antimetabolites, mimicking natural nucleotides to interfere with essential cellular processes, such as thymidylate synthesis, nucleotide reduction, DNA replication, and RNA synthesis. Examples of therapeutic deoxyribonucleoside analogues include the use of zidovudine (3’-azidothymidine) against Human Immunodeficiency Virus (HIV) (Cihlar and Ray, 2010), acyclovir (acycloguanosine) against Herpes Simplex Virus (HSV) (Clercq, 2007), as well as gemcitabine (2’,2’-difluoro-2’-deoxycytidine) (Hertel et al., 1990), cytarabine (cytosine arabinoside) (Cohen, 1976) or floxuridine (5-fluorodeoxyuridine, FdU) (Burchenal et al., 1959) to stop the proliferation of cancer cells (Mathews, 2012).

When cancer cells take up FdU, they first convert it to 5-fluorodeoxyuridine monophosphate (FdUMP) and then further phosphorylate it to become 5-fluorodeoxyuridine triphosphate (FdUTP). FdUMP competitively inhibits thymidylate synthase (TS), thus hindering the methylation of deoxyuridine monophosphate (dUMP) to obtain deoxythymidine monophosphate (dTMP). In this way, FdUMP depletes cells of deoxythymidine triphosphate (dTTP), disabling DNA synthesis. Moreover, FdUTP is incorporated into nascent DNA and induces DNA damage. Interfering with DNA repair and/or DNA damage signalling, such as base excision repair, ATM/ATR kinase activity, or poly(ADP-Ribose) polymerase, augments the cytotoxic effect of FdU (Geng et al., 2011; Huehls et al., 2016; Huehls et al., 2011;

The use of damaging nucleoside analogues against *P. falciparum* has been precluded so far, due to the parasite’s unique biology. Unlike human cells, *P. falciparum* lacks nucleoside kinases, the enzymes necessary for phosphorylating nucleosides into active nucleotide forms. The parasites take up purines mostly as hypoxanthine from the environment, followed by synthesis of inosine monophosphate as a precursor of purine nucleotides. In the case of pyrimidines, parasites synthesize uridine monophosphate from carbamoyl phosphate and aspartate, through orotate as an intermediate metabolite, but they cannot convert exogenous uridine or cytidine into nucleotides and nucleic acids (Babai et al., 2022; Büngener and Nielsen, 1967, 1968, 1969; Sherman, 1979). As a result, a fluorinated derivative of orotate, 5’-fluoroorotate / 5’-fluoroorotic acid (5-FOA), has been used to interfere with the proliferation of *Plasmodium* (Gómez and Rathod, 1990; Rathod et al., 1989; Rathod et al., 1992), but nucleoside analogues remain largely ineffective. For instance, the IC_50_ of FdU in *P. falciparum* was found to be greater than 30 µM (Queen et al., 1990). Conversely, direct administration of nucleotides, such as nucleoside triphosphates, is hampered by poor membrane permeability, which largely precludes uptake by any cell or infectious organism.

To overcome these challenges, cell-permeable nucleotide derivatives have emerged as a promising alternative (Meier, 2017). Nucleoside triphosphates were rendered lipophilic through the bioreversible modification of the γ-phosphate, by the addition of acyloxybenzyl (Gollnest et al., 2015; Jia et al., 2020), alkyl (Jia et al., 2023a) or mixed non-symmetric acyloxybenzyl-alkyl residues (Zhao et al., 2020). This was first exemplified by modifying the anti-HIV nucleoside stavudine (d4T), followed by 3’-fluoro-3’-deoxythymidine (FLT) (Weising et al., 2022) and various other nucleoside analogues. The basic idea behind these so-called Tri*PPP*ro-compounds is that the masking group(s) is/are cleaved enzymatically within the masking group, which triggers a second chemical cleavage yielding the charged phosphate moiety (Gollnest et al., 2016). In contrast, the γ-bis-alkylated triphosphate derivatives are slowly hydrolysed chemically at pH 7.3 (Jia et al., 2023b). In all cases, the compounds proved highly effective against the propagation of HIV. These previous studies suggest that similar compounds might also bypass the lack of nucleoside kinases in *P. falciparum* by enabling direct delivery of the active nucleotides into infected RBCs.

These considerations prompted us to explore the potential of FdUTP derivatives with improved cell permeability (cell-permeable FdUTP, cpFdUTP) to inhibit *P. falciparum* proliferation. We observed that cpFdUTP effectively eliminates *P. falciparum* from *in vitro* cultures by arresting parasite development at the trophozoite-to-schizont stage transition. Mechanistic studies using a *P. falciparum* nuclear cycle sensor line (Klaus et al., 2022) revealed that cpFdUTP prolongs nuclear accumulation of the DNA replication protein proliferating cell nuclear antigen 1 (PCNA1), strongly suggesting stalled DNA replication. Importantly, while cpFdUTP also inhibited the proliferation of human cells, its toxicity was completely reversed by supplementation with thymidine or cell-permeable thymidine triphosphate (cpdTTP). In contrast, these rescue agents had no impact on the anti-parasitic efficacy of cpFdUTP, highlighting a potential therapeutic window for selective parasite targeting. This study not only underscores the potential of cpFdUTP as a novel antimalarial agent but also introduces a strategic approach to mitigate host toxicity through the co-administration of thymidine. In summary, our findings pave the way for further investigation into the clinical application of cell-permeable nucleotide derivatives for malaria treatment.

## RESULTS

### cpFdUTP inhibits the proliferation of *P. falciparum in vitro*

To investigate whether cpFdUTP is active against *P. falciparum*, we propagated the strain 3D7 in human RBCs under standard conditions, while adding four different cell-permeable derivatives of FdUTP to their media at a range of concentrations. All four compounds had two lipophilic masks at the γ-phosphate (Figure 1A). We then assessed the proliferation of the parasites by SYBR Green I staining of the parasite’s DNA. Strikingly, cpFdUTP-01 strongly interfered with the proliferation of the parasites, to a far greater extent than the nucleoside analogue FdU (Figure 1B and Suppl. Figure 1). Moreover, all four cpFdUTPs (01, 02, 03 and 04) suppressed the proliferation of *P. falciparum*, with IC_50_ values of 1.871 µM, 3.168 µM, 3.318 µM and 2.145 µM, respectively (Figure 1C and Suppl. Figure 2), which are more than ten times lower than the reported IC_50_ of FdU (Queen et al., 1990). When assessing the numbers of metabolically active parasites through an assay that quantifies *P. falciparum*-specific lactate dehydrogenase (LDH) activity, all four compounds were again found active against the parasites (Suppl. Figure 3). Compound FdUTP-01 (γ-Bis(4-nonanoyloxybenzyl)-2‘-deoxy-5-fluoro-uridine-5’-triphosphate), carrying two acyloxybenzyl moieties at the γ-phosphate, was the most effective one, which prompted us to use it for mechanistic investigations. Diminished parasitemia in response to FdUTP-01 was also observed by Giemsa staining (Suppl. Figure 4, A and B). In conclusion, different cpFdUTP derivatives are capable of inhibiting *P. falciparum* growth at low micromolar concentrations.

**Figure 1:**
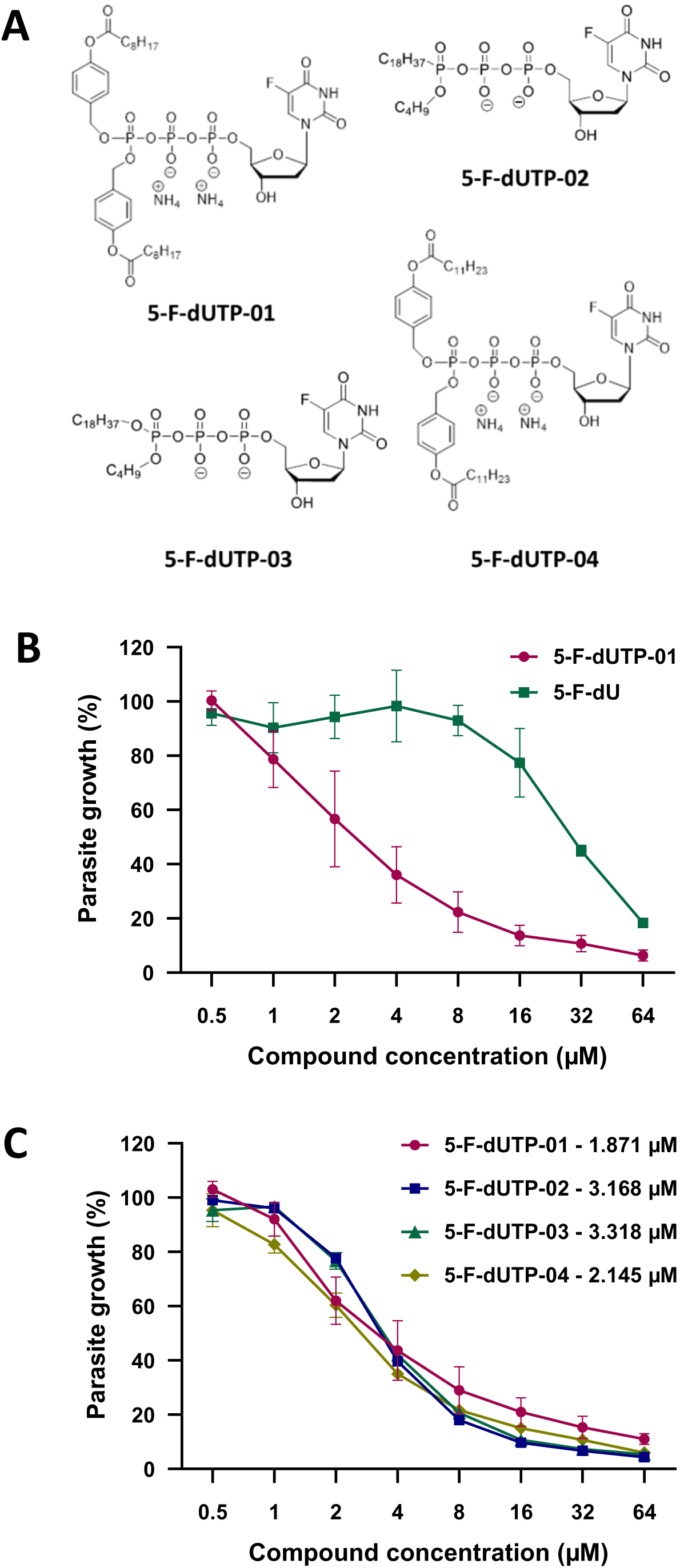
Impact of cpFdUTP on the proliferation of *P. falciparum* 3D7 *in vitro*. **A.** Structures of lipid-conjugated, cell-permeable FdUTP derivatives. FdUTP was conjugated to different lipophilic moieties, each at the γ-phosphate. 5-F-dUTP-01: γ-Bis(4-nonanoyloxybenzyl)-2‘-deoxy-5-fluoro-uridine-5’-triphosphate; 5-F-dUTP-02: γ-(Butyl)octadecyl-2‘-deoxy-5-fluoro-uridine-5‘-triphosphate; 5-F-dUTP-03: γ-(Butyl)octadecyl-2‘-deoxy-5-fluoro-uridine-5‘-triphosphonate; 5-F-dUTP-04: γ-Bis(4-dodecanoyloxybenzyl)-2‘-deoxy-5-fluoro-uridine-5’-triphosphate **B.** Impact of 5-F-dUTP-01 in comparison to 5-F-dU on *P. falciparum*. *P. falciparum* parasites, strain 3D7, were cultured asynchronously in RBCs under standard conditions, and their abundance was determined by SYBR Green I staining of *P. falciparum* DNA, followed by evaluation by automated fluorimetry. The signal obtained from non-treated parasites was used as a reference, and the reduction of the SYBR Green I positive fraction of RBCs upon addition of each drug was determined as a percentage for the indicated drug concentrations. Suppl. Fig. 1, A-C displays single replicates and raw data. Suppl. Fig. 4 shows analogous experiments using Giemsa stains as readouts. **C.** Impaired proliferation of *P. falciparum* in the presence of the FdUTP conjugates. Parasites were cultivated and treated with the FdUTP derivatives as in (**B.**). The IC_50_ value of each FdUTP conjugate is indicated. Cf. Suppl. Fig. 2, A and B, for single replicates and raw data. Similar assays using LDH activity as a readout are provided in Suppl. Fig. 3.

### cpFdUTP sustainably eliminates *P. falciparum* from culture even after compound withdrawal after 48 hours

Next, we asked whether pulsed treatment with cpFdUTP (compound FdUTP-01) was still capable of eliminating the parasites in culture. We treated asynchronously growing cultures with cpFdUTP for 24 or 48 hours and subsequently let the parasites grow in compound-free media. When the compound was removed at 24 hours, the growth of the parasites was quickly restored (Figure 2A, Suppl. Fig. 5 and 6). In contrast, treatment for 48 hours led to diminished parasitemia even after drug washout, and the parasites were hardly detectable 96 hours after removing the compound (Figure 2B, Suppl. Fig. 5 and 7). In conclusion, cpFdUTP likely acts cytocidal, rather than cytostatic, and the parasites can be eliminated within one 48-hour developmental cycle, with little if any viable parasites emerging during the subsequent days.

**Figure 2:**
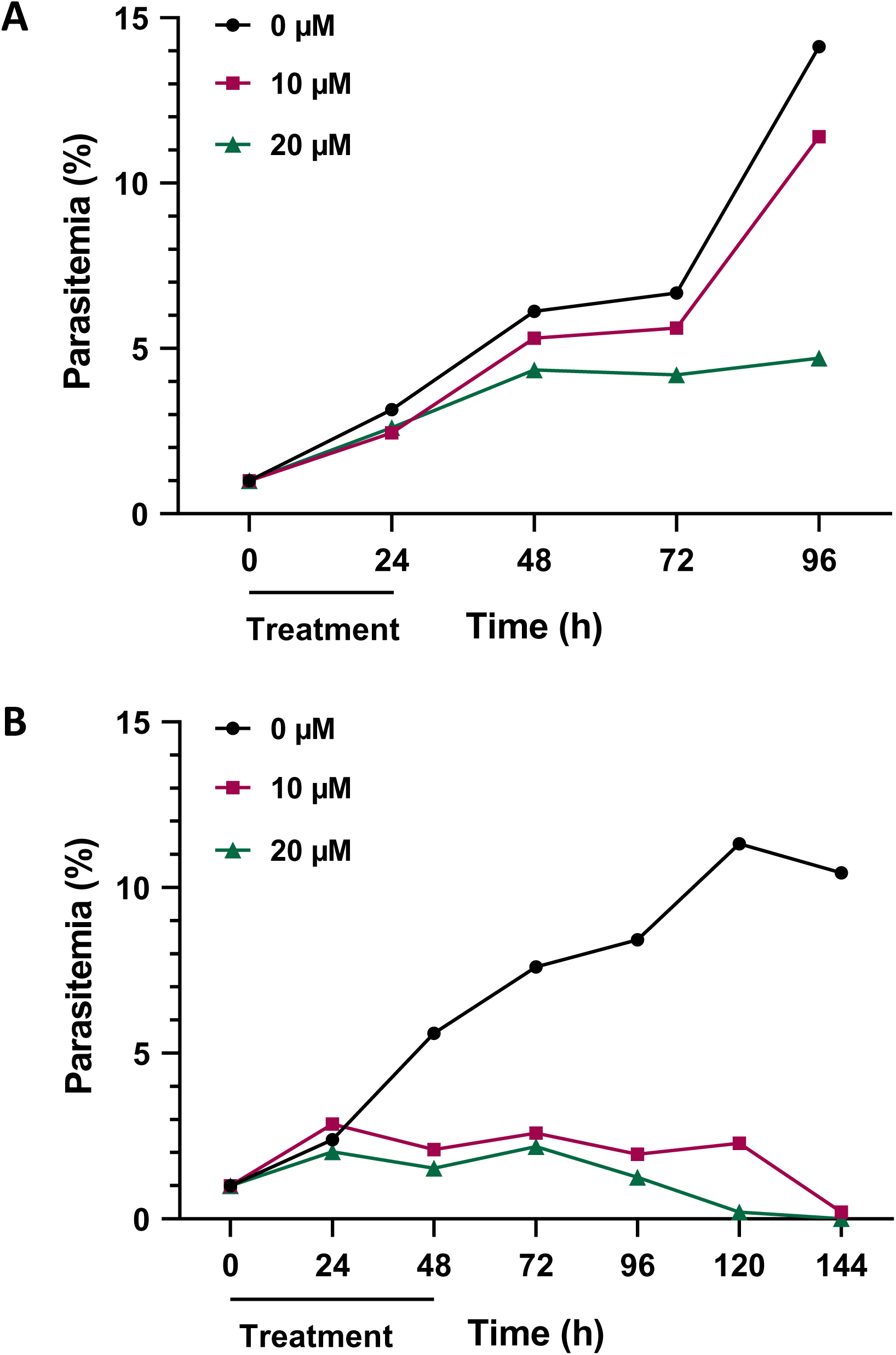
Sustainability of *P. falciparum* growth inhibition by cpFdUTP. Mixed-stage *P. falciparum* 3D7 parasites were incubated with cpFdUTP at the indicated concentrations for 24 (**A.**) or 48 (**B.**) hours, followed by further incubation of the culture. Parasitemia was determined by Giemsa staining. A total of approx. 3000 parasites were counted from three biological replicates, and a representative replicate is shown here. Parasites quickly recovered after 24 but not 48 hours of incubation with the compound. Suppl. Fig. 5 displays the other two biological replicates, and raw data are presented in Suppl. Fig. 6 and 7.

### cpFdUTP is toxic to human cells, but its toxicity is abolished by supplementation with thymidine or a cell-permeable dTTP derivative (cpdTTP)

To assess a potential therapeutic window, we tested the toxicity of cpFdUTP against human cells. Cultured Retinal Pigment Epithelia (RPE) cells, which are non-transformed, as well as lung adenocarcinoma-derived H1299 cells, were incubated with cpFdUTP, and cell confluence was measured over time. Both cell lines were susceptible to growth inhibition by the compound, even at concentrations down to 1 µM (Figure 3, A and B). As such, this would likely preclude the therapeutic use of cpFdUTP against malaria in a clinical setting. However, we reasoned that cpFdUTP might provide much of its cytotoxic effect through the inhibition of human thymidylate synthase (TS). If true, an external source of deoxythymidine nucleotides would compensate for the lack of TS activity. We therefore prepared a cell-permeable derivative of dTTP by adding two acyloxybenzyl moieties (cpdTTP; γ-bis(4-decanoyloxybenzyl)-2’-deoxy-thymidine-5’-triphosphate; Figure 3C). As such, cpdTTP had no observable effect on cell proliferation. However, when used in combination, cpdTTP completely rescued H1299 and RPE cells from the cytotoxicity of cpFdUTP. This was true even when cpdTTP was used at 5- to 10-fold lower concentrations than cpFdUTP (Figure 3, D, E and F). Finally, we reasoned that mammalian cells contain nucleoside kinases and ribonucleotide reductase to synthesize deoxythymidine nucleotides from thymidine, a cell-permeable nucleoside. And indeed, non-modified thymidine was also capable of completely rescuing both H1299 and RPE cells from cpFdUTP (Figure 3, G, H and I). These observations argue that, despite the cytotoxicity of cpFdUTP, the addition of thymidine or cpdTTP allows human cells to survive without detectable damage, possibly constituting a therapeutic window.

**Figure 3:**
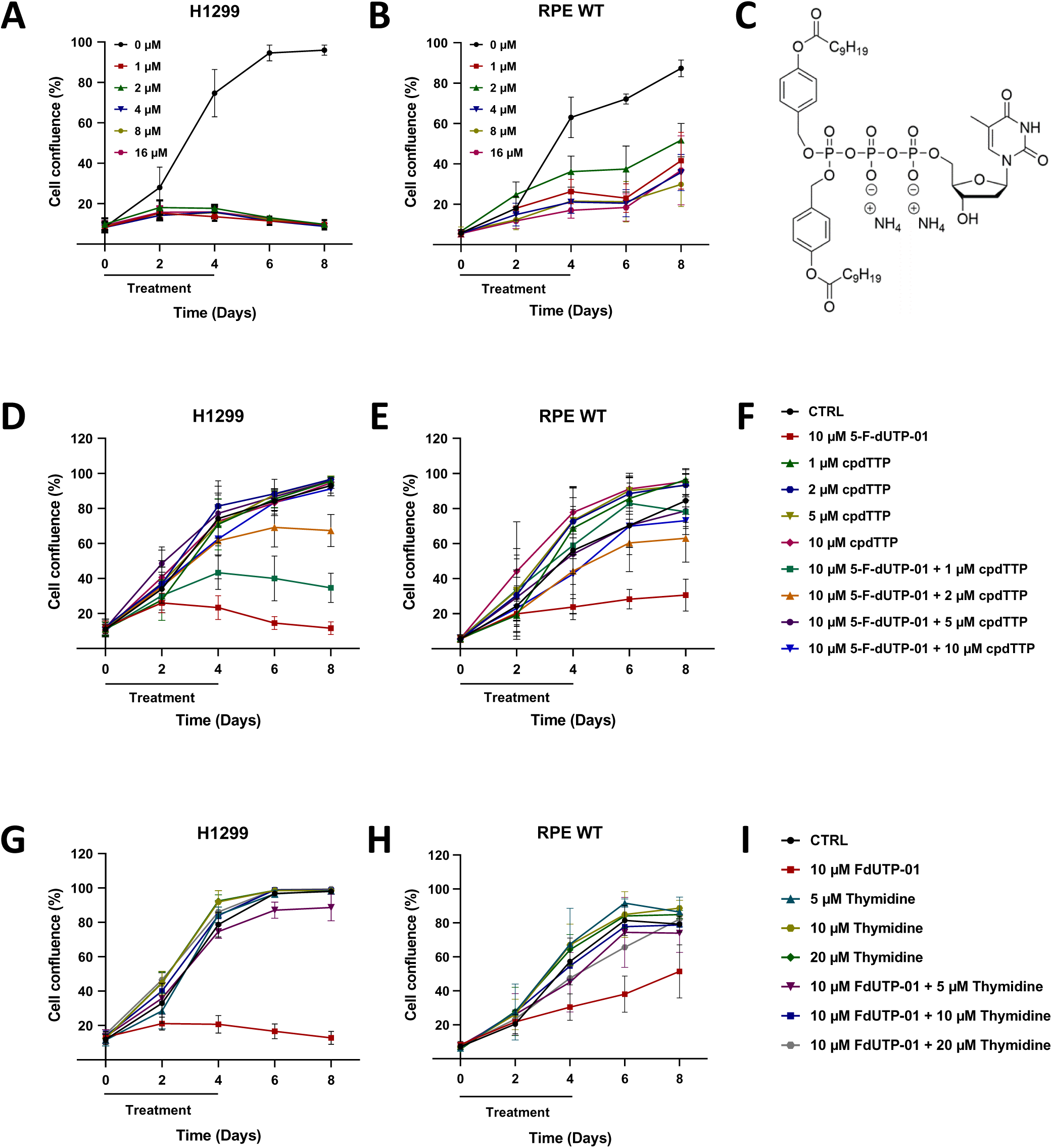
Reversible toxicity of cpFdUTP on human cells. Human cells were incubated with cpFdUTP at increasing concentrations as indicated. Cell confluence was measured by automated microscopy during four days of incubation and four subsequent days after compound withdrawal. **A.** The effect of cpFdUTP on H1299 cells, derived from human lung carcinoma, was studied. **B.** The response of RPE cells, a non-transformed cell line from human retinal pigment epithelia, to cpFdUTP was investigated. **C.** Structure of a cell-permeable (cp) derivative of dTTP with lipid-like modifications at the γ-phosphate. The impact of cpdTTP on the intraerythrocytic proliferation of *P. falciparum* is shown in Suppl. Fig. 8. **D.** cpFdUTP was combined with cpdTTP, which abolished the toxicity towards H1299 cells. **E.** In RPE WT cells, cpdTTP alleviated the toxicity of cpFdUTP. **F.** Legend to (**D.**) and (**E.**) **G.** Thymidine alleviates the toxicity of cpFdUTP on cpFdUTP-treated H1299 cells. **H.** Thymidine rescues the proliferation of RPE cells treated with cpFdUTP. **I.** Legend to (**G.**) and (**H.**)

### Neither thymidine, nor cpdTTP can compromise the anti-parasitic activity of cpFdUTP

Next, we tested whether thymidine or cpdTTP might also impair the therapeutic impact of cpFdUTP on *P. falciparum*. When adding thymidine or cpdTTP to cultured parasites, we observed reduced growth only at concentrations over 50 µM (Suppl. Figure 8, A and B). Most notably, however, when we combined thymidine or cpdTTP with cpFdUTP to treat *P. falciparum* parasites, we did not observe any rescue effect (Figure 4, A and B, and Suppl. Fig. 9 and 10). These findings imply that the combination of cpFdUTP with thymidine or cpdTTP could open a therapeutic window that allows the elimination of *P. falciparum* while causing no harm to human cells. Moreover, these results argue that cpFdUTP interferes with the growth of human cells and *P. falciparum* by different predominant mechanisms. We propose that the proliferation of human cells is hindered mostly through TS inhibition, e.g. by the cpFdUTP metabolite FdUMP. In contrast, cpFdUTP appears to suppress the growth of *P. falciparum* without critical depletion of thymidine deoxyribonucleotides, which is in agreement with abundant dTTP levels in the parasites (Babai et al., 2022). Instead, cpFdUTP might hinder S phase progression, e.g. through incorporation into nascent DNA.

**Figure 4:**
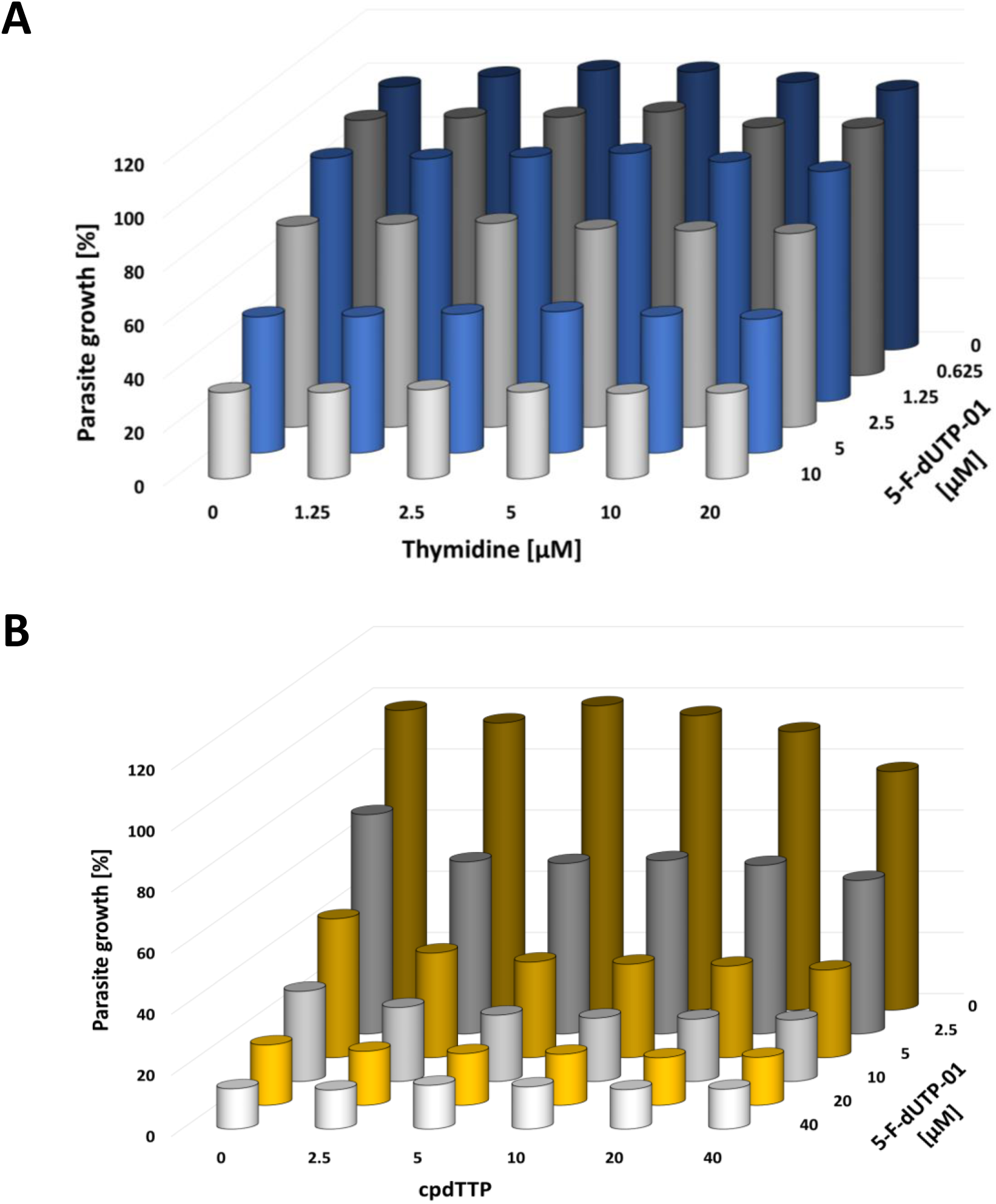
Failure of thymidine or cpdTTP to rescue *P. falciparum* growth upon cpFdUTP treatment. **A.** *P. falciparum* cultures were incubated with cpFdUTP, and the proliferation of *P. falciparum* was assessed by SYBR Green staining as in Figure 1B. cpFdUTP was combined with either thymidine (**A.**) or cpdTTP (**B.**), but neither of the compounds rescued *P. falciparum* from the deleterious impact of cpFdUTP. Suppl. Fig. 9 and 10 display the individual 2D bar graphs for each concentration of thymidine and cpdTTP, respectively.

### cpFdUTP, when applied to *P. falciparum* ring stages, prevents the subsequent round of invasion

To gain further insight into the anti-parasitic activity of cpFdUTP, we synchronized *P. falciparum* by sorbitol treatment (Lambros and Vanderberg, 1979) and added the compound to a ring-stage culture (Figure 5A). This prevented the increase in parasitemia over the subsequent 72 hours (Figure 5B). It also precluded the re-appearance of ring stages at 48 hours of treatment (Figure 5C), compatible with the view that no infectious parasites can form in the presence of cpFdUTP. Instead, we observed the accumulation of parasites with abnormal morphologies (Figure 5C, D). Similar patterns were observed when the parasites were treated with 5-fluoroorotic acid (5-FOA), an established inhibitor of thymidylate synthase (Figure 5D). In conclusion, cpFdUTP treatment interfered with the development of late parasite stages, consistent with a perturbation of DNA metabolism, thus precluding the release of infectious parasites and the subsequent round of invasion.

**Figure 5:**
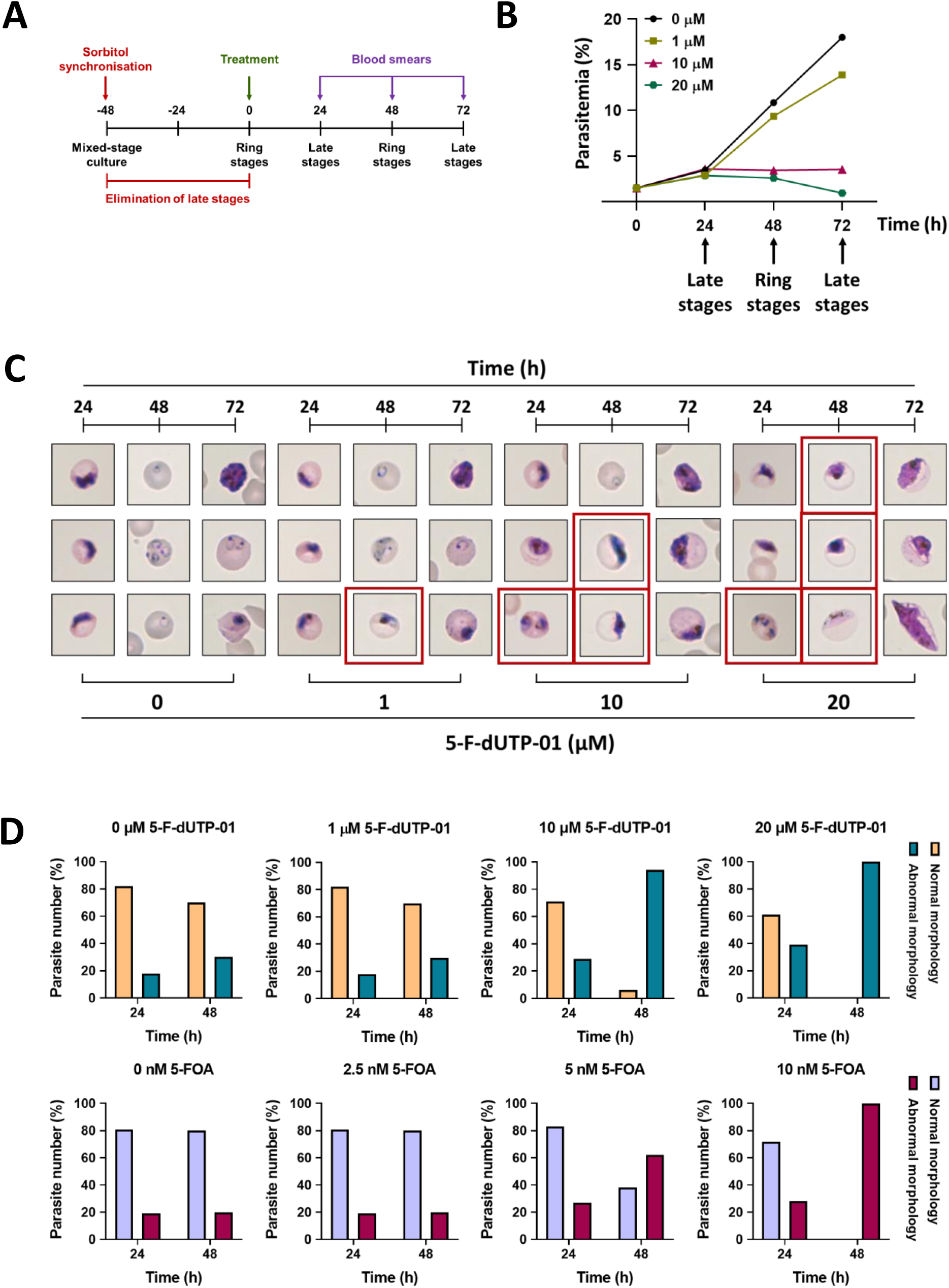
Arrest of *P. falciparum* at the trophozoite stage by cpFdUTP. **A.** Scheme indicating the experimental setup and synchronization of *P. falciparum*. Sorbitol treatment eliminated late-stage parasites (trophozoites and schizonts), leaving ring stages (again) at 48 hours post treatment. At this time, treatment with cpFdUTP was started, followed by Giemsa staining of samples at 24, 48 and 72 hours of incubation with cpFdUTP, as well as of samples from non-treated parasites. **B.** Reduced parasitemia in response to cpFdUTP. The expected parasite stages are indicated below the graph **C.** Representative images of *P. falciparum*-infected RBCs upon cpFdUTP treatment at the indicated concentrations and durations. Ring stages were no longer observed at higher cpFdUTP concentrations. Parasites with abnormal morphologies are framed in red. At higher cpFdUTP concentrations, abnormal parasites appear already at 24 hours of treatment and represent the majority of parasites after 48 hours. After 72 hours of treatment, no asexual *P. falciparum* stages could be detected at 20 µM cpFdUTP, with only gametocytes being present in the culture. **D.** Quantifications of parasite morphologies upon synchronization and subsequent treatment with cpFdUTP or 5-fluoroorotic acid (5-FOA) for comparison. A total of 100 parasites from 2 biological replicates were quantified.

### cpFdUTP hinders S phase progression in *P. falciparum*

Finally, we asked whether cpFdUTP interferes with the replication of *P. falciparum* DNA. To elucidate this, we took advantage of a previously established *P. falciparum* nuclear cycle sensor line, which is based on Green Fluorescent Protein (GFP)-tagged Proliferating Cell Nuclear Antigen 1 (PCNA1). *P. falciparum* PCNA1 is part of the DNA replication fork. It transiently accumulates only in those nuclei of a multinucleated schizont that replicate their DNA. Thus, nuclear accumulation of PCNA1::GFP can serve as a proxy for S phase progression. This can be quantitatively assessed as nuclear accumulation of PCNA1::GFP. S phase coincides with an increase in peak GFP signal intensity, which decreases again as DNA replication concludes (Klaus et al., 2022). Consistently, we found that treatment with aphidicolin, which blocks DNA replication in *P. falciparum* (Machado et al., 2023), led to a prolonged nuclear accumulation of PCNA1::GFP, as compared to untreated parasites (Figure 6A-D). Strikingly, the addition of cpFdUTP led to a stronger increase in the peak GFP signal intensity, compared to aphidicolin, and an extensively prolonged nuclear accumulation of *P. falciparum* PCNA1 (Figure 6E, F; Supplemental Movie M1). Since cpdTTP did not overcome the anti-parasitic activity of cpFdUTP (Figure 4B and Suppl. Fig. 10), we propose that cpFdUTP hinders the progression of DNA synthesis, perhaps through incorporation into nascent DNA strands. In this scenario, stalled replication forks and Okazaki fragments will remain, all of which have PCNA1 bound, extending the nuclear accumulation of PCNA1. In conclusion, these observations strongly suggest that cpFdUTP hinders the progression of *P. falciparum* through S phase.

**Figure 6:**
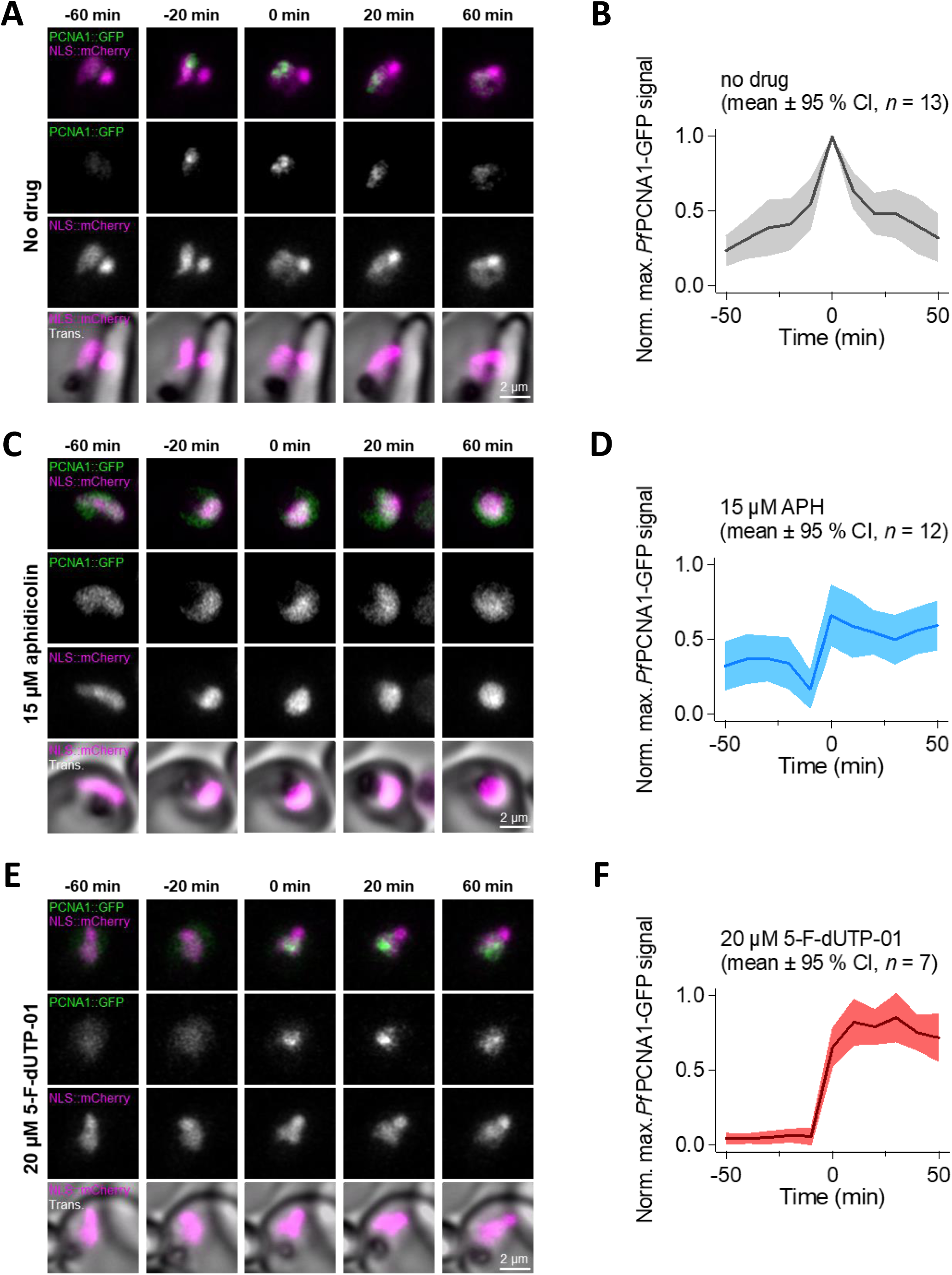
Prolonged nuclear accumulation of PCNA1 in cpFdUTP-treated *P. falciparum* indicates stalled DNA replication. **A.** Live-cell imaging of early schizonts that episomally express PCNA1::GFP and 3xNLS::mCherry, which marks the nuclei. Note, nuclear accumulation of PCNA1::GFP occurs during DNA replication in a given nucleus (Klaus et al., 2022). **B.** Maximal PCNA1::GFP signal over time of untreated parasites. Note, nuclear accumulation of PCNA1::GFP resulted in a peak of the maximal pixel intensity, which was aligned to time = 0 min. The GFP signal was normalized to the weakest (= 0) and the strongest (= 1) maximal pixel value from the beginning of imaging to 6 hours after nuclear accumulation of PCNA1::GFP. **C.** Live-cell imaging of early schizonts as in (**A.**), treated with 15 µM aphidicolin (APH). **D.** Maximal PCNA1::GFP signal over time of schizonts as in (**B.**), treated with 15 µM APH. **E.** Live-cell imaging of early schizonts as in (**A.**), treated with 20 µM 5-F-dUTP-01. Suppl. Movie M1 is a time-lapse video generated from the images of parasites treated with 20 µM 5-F-dUTP-01. **F.** Maximal PCNA1::GFP signal over time of schizonts as in (**B.**), treated with 20 µM 5-F-dUTP-01. Scale bar = 2 µm

## DISCUSSION

Our results indicate anti-parasitic activity of a lipid-like modified nucleotide analogue, cpFdUTP, against the human malaria parasite *P. falciparum*. Unlike nucleoside analogues, this compound strongly interferes with the proliferation of the parasite. The toxicity of the compound in human cells is abolished by the addition of cpdTTP or thymidine. However, neither thymidine, nor cpdTTP diminished the therapeutic impact of cpFdUTP on *P. falciparum*, which opens a therapeutic window towards a novel principle of antimalarial therapy.

Nucleoside analogues have been in use for decades against viruses and cancer. However, none of them showed activity against *Plasmodium*, in full agreement with the absence of nucleoside kinases in these parasites (McCormick et al., 1974; Merrick, 2015). Thus, only the phosphorylation of nucleosides (to become nucleotides) can make this class of drugs active against the parasite. Since plain nucleotides are too polar to permeate a lipid membrane, lipophilic modifications are key to their use as drug candidates (Gollnest et al., 2015). Such modifications raise the perspective of converting an entire class of antiviral and anticancer drugs to become novel antimalarials.

The need for new antimalarial drugs is largely due to the development of resistance towards existing ones. Even the currently most successful class of antimalarial drugs, artemisinin and its derivatives, has given rise to less susceptible mutants (Rosenthal et al., 2024). Although our study does not include analyses of long-term resistance development towards cpFdUTP, the proposed mechanism of action, i.e. incorporation into nascent DNA, suggests that resistance will be difficult to achieve by single mutations. Mammalian cells also incorporate fluorinated uridine derivatives through multiple machineries, e.g. DNA polymerases, and there is no known mutation in these polymerases that would decrease the incorporation of FdUTP vs. dTTP. Thus, DNA-incorporated nucleotide analogues might be developable as a class of antimalarial drug candidates with a relatively small risk of resistance development.

The development of membrane-permeable nucleotides raises the perspective of using additional nucleotides with similar modifications against *Plasmodium*. In the case of viruses, chain-terminating compounds are used against viral DNA polymerases (Seley-Radtke and Yates, 2018; Yates and Seley-Radtke, 2019). These nucleotides typically lack a 3’ hydroxyl group that would be required for the formation of a phosphodiester bond with the subsequent nucleotide. However, immediate chain terminators can be excised through the proofreading activity of most non-viral DNA polymerases. This proofreading activity was also found in *Plasmodium* DNA polymerases in the nucleus (Kümpornsin et al., 2023), the apicoplast (Sharma et al., 2020; Wingert et al., 2013) and mitochondria (Chavalitshewinkoon-Petmitr et al., 2000). Therefore, this activity might render *Plasmodium* resistant against immediate chain terminators. However, delayed chain termination by nucleoside analogues can hinder the proliferation of cancer cells upon incorporation by mammalian DNA polymerases (Galmarini et al., 2002). These compounds hinder further 3’ extension only after incorporating a few additional nucleotides, limiting the efficacy of the proofreading 3’-5’ exonucleoase activity. Examples include the cytidine analogues cytosine arabinoside (Ara-C) or gemcitabine. Converting similar nucleoside analogues to cell-permeable nucleotides might make them active against malaria-causing parasites, especially when combined with the corresponding natural nucleoside to protect the host cells.

Taken together, applying lipid-like modified nucleotide analogues towards *P. falciparum* has the potential of establishing a hitherto unexploited therapeutic principle to treat malaria.

## Supporting information

Supplemental Movie M1: Prolonged nuclear localization of PCNA1 after exposure to cpFdUTP. Corresponding to Figure 6E.

## ACKNOWLEDGEMENTS

During this work, VN was a scholar of the International Max Planck Research School for Molecular Biology, part of the GGNB graduate school.

The authors are grateful to the Infectious Diseases Imaging Platform (www.idip-heidelberg.org) and the *Plasmodium* database PlasmoDB (www.plasmodb.org), which facilitated this work. This study was supported by the Health + Life Science Alliance Heidelberg Mannheim, receiving state funding approved by the State Parliament of Baden-Württemberg, and the German Research Foundation - project number 240245660 - SFB 1129 to MG.

## AUTHOR CONTRIBUTIONS

Conceptualization: MG, CM, MD

Experimentation: VN, KL, M-EB, BG, PB, MV, MM

Supervision: MG, CM, MD

Manuscript, initial draft: MD

Manuscript, correction and finalization: all authors

## MATERIALS AND METHODS

### Cell culture

The human non-small cell lung cancer cell line H1299 was maintained in Dulbecco’s Modified Eagle’s Medium (DMEM, Thermo Fisher Scientific) supplemented with 10% fetal bovine serum (FBS, Anprotec), 2 mM glutamine (Thermo Fisher Scientific), 50 U/mL penicillin, 50 μg/mL streptomycin (Gibco), 10 μg/mL ciprofloxacin (Fresenius Kabi) and 2 μg/mL tetracycline (Sigma Aldrich). The wildtype non-tumorigenic human retinal pigment epithelial cell line RPE was cultured in DMEM GlutaMAX (Gibco) supplemented with 10% FBS, penicillin, streptomycin, ciprofloxacin and 50 μg/mL hygromycin B (Thermo Fisher Scientific) for hTERT maintenance. All cell lines were maintained at 37°C with 5% CO_2_, and were routinely tested for *Mycoplasma* contamination.

### Confluence measurements of human cells

Cells were plated on translucent 12-well plates (Corning Costar) and continuously treated with the indicated compounds starting on the day after seeding. The cells were treated for a total of 96 hours, during which the treatment was refreshed every 48 hours. Following that, cells were allowed to grow in compound-free media for another 96 hours, during which the medium was refreshed every 48 hours. On each day of treatment, cell confluence was measured and analysed using an automated imaging cytometer (Celigo, Nexcelom Bioscience). GraphPad Prism 8.0 was used for data analysis and graph generation.

### *In vitro* cultivation of *P. falciparum* 3D7 in human RBCs

*P. falciparum* parasites (strain 3D7) were obtained from Hosam Shams-Eldin (Marburg University, Germany). Asexual blood-stage parasites were cultivated in AB-positive or AB-negative human erythrocytes (German Red Cross blood bank, DRK-NSTOB) at approx. 4% hematocrit in homemade RPMI-1640 medium (Gibco) supplemented with 5.95 g/L HEPES (Carl Roth), 50 mg/L hypoxanthine (Carl Roth), 0.2% NaHCO_3_ (Carl Roth), 0.5% AlbuMAX-I (Gibco) and 12.5 µg/mL gentamycin (Sigma Aldrich). Cultures were maintained at 37°C in hypoxia chambers (STEMCELL Technologies) under an atmosphere consisting of 95% N_2_, 5% O_2_ and 5% CO_2_.

### Compound preparation

The synthesis of Tri*PPP*ro-NTP prodrugs followed our previously reported protocols (Gollnest et al., 2015; Jia et al., 2023a).

Under nitrogen as inert gas, the corresponding *H*-phosphonate (1.0 equiv.) was dissolved/suspended together with *N*-chlorosuccinimide (2.0 equiv.) in acetonitrile and stirred at 60°C for 1 hour. Monobasic tetra-n-butylammonium phosphate (3.0 equiv., 0.4 mol/L in acetonitrile) was added, and the reaction was stirred at 60°C for 1 hour. After all volatiles were removed under reduced pressure, the residue was taken up in dichloromethane and the organic phase was washed consecutively with 1 M aq. ammonium acetate (1 mol/L) and water, then dried over Na_2_SO_4_ (phase separation via centrifugation). Finally, all volatiles were removed under reduced pressure and the obtained pyrophosphate was used without further purification (storage was carried out under nitrogen as an inert gas and at −20°C).

The pyrophosphate (1.0 equiv.) was dissolved in acetonitrile, and a solution of triethylamine (16 equiv.) and trifluoroacetic anhydride (10 equiv.) was added dropwise. After stirring for 10 minutes at room temperature, all volatiles were removed under reduced pressure, the residue was taken up in DMF, and triethylamine (10 equiv.) and 1-methylimidazole (6.0 equiv.) were added. After stirring for 10 minutes at room temperature, the appropriate monophosphate (0.5 - 0.7 equiv.), dissolved in DMF, was added. The reaction was stirred until complete conversion (RP_18_ HPLC monitoring) at room temperature, after which all volatiles were removed under reduced pressure. The residue was purified on RP_18_ silica gel using automated flash chromatography, followed by ion exchange to ammonia (DOWEX-NH_4_^+^), and a final purification on RP_18_ silica gel using automated flash chromatography. Subsequently, the compounds were characterized by NMR analysis as follows.

#### 5-F-dUTP-01

1H-NMR: (600 MHz, MeOH-*d*_4_): δ [ppm] = 8.05 (d, J = 6.6 Hz, 1H), 7.42 - 7.37 (m, 4H), 7.05 - 7.01 (m, 4H), 6.25 (td, J = 6.9 Hz, J = 1.7 Hz, 1H), 5.16 (d, J = 8.9 Hz, 4H), 4.56 (dt, J = 5.9 Hz, J = 3.2 Hz, 1H), 4.25 (ddd, J = 11.3 Hz, J = 6.4 Hz, J = 3.2 Hz, 1H), 4.20 - 4.14 (m, 1H), 4.02 - 3.99 (m, 1H), 2.56 (t, J = 7.4 Hz, 4H), 2.25 - 2.16 (m, 2H), 1.72 (p, J = 6.9 Hz, 4H), 1.45 - 1.38 (m, 4H), 1.38 - 1.25 (m, 16H), 0.90 (t, J = 6.9 Hz, 6H). **^31^*P*-NMR:** (162 MHz, MeOH-*d*_4_): δ [ppm] = −11.67 (d, J = 19.9 Hz), −13.30 (d, J = 17.8 Hz), −23.77 (t, J = 18.8 Hz). **^19^*F*-NMR:** (162 MHz, MeOH-*d*_4_): δ [ppm] = −167.3. **HRMS** (ESI^-^): *m/z* = calcd.: 977.2809 [M-H]^-^, found: 977.2820 [M-H]^-^.

#### 5-F-dUTP-02

1H-NMR: (600 MHz, MeOH-*d*_4_): δ [ppm] = 8.11 (d, J = 6.6 Hz, 1H), 6.28 (ddd, J = 7.6 Hz, J = 5.9 Hz, J = 1.8 Hz, 1H), 4.58 (dt, J = 6.0 Hz, J = 3.1 Hz, 1H), 4.23 - 4.12 (m, 4H), 4.03 (q, J = 2.9 Hz, 1H), 2.31 - 2.22 (m, 2H), 2.07 - 1.99 (m, 2H), 1.73 - 1.60 (m, 4H), 1.48 - 1.38 (m, 4H), 1.36- 1.24 (m, 28H), 0.95 (t, J = 7.4 Hz, 3H), 0.90 (t, J = 7.0 Hz, 3H). **^31^*P*-NMR:** (162 MHz, MeOH-*d*_4_): δ [ppm] = 24.07 (d, J = 23.2 Hz), −11.7 (d, J = 19.3 Hz), −23.8 (t, J = 21.6 Hz). **^19^*F*-NMR:** (162 MHz, MeOH-*d*_4_): δ [ppm] = −167.4. **HRMS** (ESI^-^): *m/z* = calcd.: 777.3063 [M-H]^-^, found: 777.3082 [M-H]^-^.

#### 5-F-dUTP-03

1H-NMR: (600 MHz, MeOH-*d*_4_): δ [ppm] = 8.09 (d, J = 6.6 Hz, 1H), 6.28 (ddd, J = 7.5 Hz, J = 6.0 Hz, J = 1.8 Hz, 1H), 4.58 (dt, J = 5.9 Hz, J = 3.1 Hz, 1H), 4.29 - 4.24 (m, 1H), 4.21 - 4.13 (m, 5H), 4.05 - 4.02 (m, 1H), 2.30 - 2.23 (m, 2H), 1.72 - 1.65 (h, J = 6.6 Hz, 4H), 1.48 - 1.37 (m, 4H), 1.36 - 1.26 (m, 28H), 0.96 (t, J = 7.4 Hz, 3H), 0.90 (t, J = 7.0 Hz, 3H). **^31^*P*-NMR:** (243 MHz, MeOH-*d*_4_): δ [ppm] = −11.9 (d, J = 18.4 Hz), −11.7 (d, J = 19.5 Hz), −24.1 (t, J = 21.6 Hz). **^19^*F*-NMR:** (565 MHz, MeOH-*d*_4_): δ [ppm] = −167.5. **HRMS** (ESI^-^): *m/z* = calcd.: 793.3012 [M-H]^-^, found: 793.2998 [M-H]^-^.

#### 5-F-dUTP-04

1H-NMR: (600 MHz, MeOH-*d*_4_): δ [ppm] = 8.06 (d, J = 6.5 Hz, 1H), 7.42 - 7.38 (m, 4H), 7.06 - 7.02 (m, 4H), 6.26 (ddd, J = 7.6 Hz, J = 6.1 Hz, J = 1.7 Hz, 1H), 5.17 (d, J = 8.3 Hz, 4H), 4.57 (dt, J = 5.9 Hz, J = 3.1 Hz, 1H), 4.27 (ddd, J = 11.3 Hz, J = 6.3 Hz, J = 3.1 Hz, 1H), 4.19 (dt, J = 11.3 Hz, J = 4.4 Hz, 1H), 4.03 - 4.00 (m, 1H), 2.57 (t, J = 7.4 Hz, 4H), 2.26 - 2.17 (m, 2H), 1.73 (p, J = 7.5 Hz, 4H), 1.46 - 1.40 (m, 4H), 1.40 - 1.26 (m, 28H), 0.90 (t, J = 7.0 Hz, 6H). **^31^*P*-NMR:** (243 MHz, MeOH-*d*_4_): δ [ppm] = −11.73 (d, J = 20.5 Hz), −13.32 (d, J = 16.8 Hz), - 23.89 (t, J = 19.1 Hz). **^19^*F*-NMR:** (565 MHz, MeOH-*d*_4_): δ [ppm] = −167.2. **HRMS** (ESI^-^): *m/z* = calcd.: 1061.3748 [M-H]^-^, found: 1061.3760 [M-H]^-^.

#### cpdTTP

**^1^*H*-NMR** (600 MHz, MeOD-*d*4): δ [ppm] = 7.80 (s, 1H), 7.40 (d, J = 8.5 Hz, 4H), 7.04 (d, J = 8.4 Hz, 4H), 6.32 (dd, J = 7.6 Hz, J = 6.04 Hz, 1H), 5.20 - 5.14 (m, 4H), 4.58 (dt, J = 6.3 Hz, J = 3.0 Hz, 1H), 4.28 - 4.20 (m, 2H), 4.02 (q, J = 2.8 Hz, 1H), 2.58 (t, J = 7.5 Hz, 4H), 2.30 - 2.23 (m, 1H), 2.19 (ddd, J = 13.5 Hz, J = 6.1 Hz, J = 3.2 Hz, 1H), 1.92 (d, J = 1.0 Hz, 3H), 1.74 (p, J = 7.6 Hz, 4H), 1.46 - 1.28 (m, 24H), 0.90 (t, J = 7.1 Hz, 6H). **^31^*P*-NMR** (243 MHz, MeOD-*d*4): δ [ppm] = −11.77 (d, J = 19.8 Hz), −13.17 (d, J = 17.2 Hz), −23.78 (t, J = 18.1 Hz).

#### HRMS

(ESI^-^): *m/z* = calcd: 1001.3372 [M-H]^-^, found: 1001.3373 [M-H]^-^.

### Compound treatment and IC_50_ value calculation

All four cell-permeable FdUTP derivatives (5-F-dUTP-01 through 5-F-dUTP-04) and the cell-permeable dTTP analogue were synthesized at the Department of Chemistry at the University of Hamburg (Germany). All stock solutions were prepared in DMSO, and were used at the indicated concentrations in the respective assays. IC_50_ values were calculated by nonlinear regression (curve fit) using GraphPad Prism.

### Giemsa staining and light microscopy for assessment of parasite morphology

Thin-blood smears were made on glass microscope slides (Labsolute), followed by fixation for 1-5 minutes in pure methanol. After air-drying, smears were incubated with Giemsa’s solution (Sigma Aldrich) diluted 1:4 in homemade Giemsa buffer [11.7 µM Na_2_HPO_4_ (Carl Roth), 4 µM KH_2_PO_4_ (Carl Roth) in ddH_2_O] for 10 minutes at room temperature. Following that, slides were thoroughly rinsed under running water and let to dry completely. For quantification and assessment of parasite morphology, a Zeiss Axio Scope.A1 microscope equipped with a Zeiss Achroplan 100x/1.25 oil immersion objective was used. Images were taken and processed using ZEN 2 (blue edition).

### Synchronization of *P. falciparum* 3D7

Parasites were synchronized by lysis of late stages using 5% D-sorbitol (Sigma Aldrich). In brief, non-infected and *P. falciparum*-infected RBCs were pelleted by centrifugation, followed by a 15-minute incubation with 5% D-sorbitol at 37°C. Atthe end, parasites were washed with complete RPMI-1640 medium by several rounds of centrifugation.

### Determination of total DNA content in *P. falciparum*-infected RBCs (SYBR Green I fluorescence-based proliferation assay)

Total DNA content of *P. falciparum* 3D7-infected erythrocytes was determined by the SYBR Green I fluorescence-based proliferation assay, following a previously described protocol with minor modifications (Leidenberger et al., 2017). In brief, 0.5% mixed-stage parasites were incubated with the compounds of interest in black 96-well plates with a transparent bottom (Falcon) for 72 hours in a hypoxic environment. Following that, parasites were lysed using homemade lysis buffer [ddH_2_O, 1x SYBR Green I (Invitrogen), 40 mM Tris-HCl pH 7.5 (Carl Roth), 10 mM EDTA pH 8.0 (Carl Roth), 0.02% Saponin (Carl Roth), 0.08% Triton X- 100 (Sigma Aldrich)], and the DNA levels were measured using Twinkle LB970 fluorescence plate reader (Berthold Technologies) at an excitation wavelength of 485 nm and an emission wavelength of 520 nm. Data analysis was conducted in Microsoft Excel, and graphs were generated using GraphPad Prism.

### Determination of *P. falciparum* LDH activity

For this assay, parasites were synchronized using 5% D-sorbitol, as described previously. Late-stage parasites at 1% parasitemia and 2% hematocrit were treated with the compounds of interest for 48 hours. An NBT solution was prepared by adding a nitroblue tetrazolium tablet (NBT, Sigma Aldrich) to an LDH buffer [ddH_2_O, 100 mM Tris-HCl pH 8.0 (Carl Roth), 0.2M sodium L-lactate (Sigma Aldrich), 0.25% Triton X-100 (Sigma Aldrich)]. The complete LDH substrate [NBT solution, 50 units/ml diaphorase from *Clostridium kluyveri* (Sigma Aldrich), 10 mg/ml 3-Acetylpyridine adenine dinucleotide (APAD, Sigma Aldrich)] was added to the parasites, followed by an incubation of 30-60 minutes at room temperature. Absorbance was measured at 650 nm using Apollo LB913 microplate spectrophotometer (Berthold Technologies), followed by data analysis in Microsoft Excel.

### Live-cell imaging of *P. falciparum* PCNA1::GFP

For live-cell imaging of *P. falciparum* PCNA1::GFP, we used a previously established cell line, which ectopically expresses PCNA1::GFP as well as mCherry with three N-terminal nuclear localization signals (NLS::mCherry), which marks the nuclei (Klaus et al., 2022). For imaging, we followed previously published protocols with minor modifications (Klaus et al., 2022).

In brief, sterile glass bottom round dishes (Ibidi) were coated with 5 mg/mL Concanavalin A (Merck) and rinsed with PBS. About 500 µL of resuspended parasite culture was washed twice with prewarmed incomplete RPMI-1640 medium and left to settle on the dish for 20 minutes at 37°C before unattached cells were washed off using incomplete RPMI until a monolayer remained. Cells were left to recover under standard culture conditions in complete RPMI until 1 hour before imaging. Following that, the media were exchanged to prewarmed phenol red-free complete RPMI imaging medium [RPMI 1640 L-Glutamine (PAN-Biotech)] supplemented with 0.5% AlbuMAX II (Thermo Fisher Scientific), 0.2 mM Hypoxanthine (cc-pro), 25 mM HEPES pH 7.3 (Sigma Aldrich), and 12.5 µg/mL gentamicin (Carl Roth) and, if needed, drugs were added. The dishes were filled with 5 ml of the imaging media and were not fully closed to allow gas exchange.

Point laser scanning confocal microscopy was performed on a Zeiss LSM900 microscope equipped with the Airyscan detector using a Plan-Apochromat 63x/1.4 oil immersion objective. Live-cell imaging was performed at 37°C and 5% CO_2_ in a humidified environment.

Images were acquired at multiple positions using an automated stage and the Definite Focus module for focus stabilization with a time-resolution of 10 minutes/stack for 12 hours. Multichannel images were acquired sequentially in the line-scanning mode using 561 nm and 488 nm diode lasers for mCherry and GFP imaging, respectively. Emission detection was configured using variable dichroic mirrors to be 570-700 for mCherry and 490-550 for GFP detection. GaAsP PMT or Airyscan detectors were used with the gain typically adjusted at 900V, offset was typically not adjusted (0%). Brightfield images were obtained from a transmitted light PMT detector. Sampling was typically Nyquist-optimised (approx. 50 nm in xy axis and 250 nm in z axis), bidirectionally with pixel dwell time between 0.7 and 1.2 μs and 2x line averaging. Subsequently, ZEN Blue 3.1 software was used for the post-2D or 3D Airyscan processing with automatically determined default Airyscan Filtering (AF) strength.

Accumulation in the nucleus of episomally expressed *P. falciparum* PCNA1::GFP was quantified as previously described (Klaus et al., 2022). In brief, we normalized the maximal GFP pixel intensity values over time, i.e. the weakest (= 0) and the strongest (= 1) maximal pixel value from the beginning of imaging to 6 hours past the nuclear accumulation of PCNA1::GFP. Maximal GFP pixel values per time point were measured from the sum of the intensities in z-projections. Cells were aligned based on the first occurrence of PCNA1::GFP nuclear accumulation, and the data were plotted using GraphPad Prism, with the accumulation event set as time zero.

## LEGENDS TO UPPLEMENTAL FIGURES

**Supplemental Figure 1:**
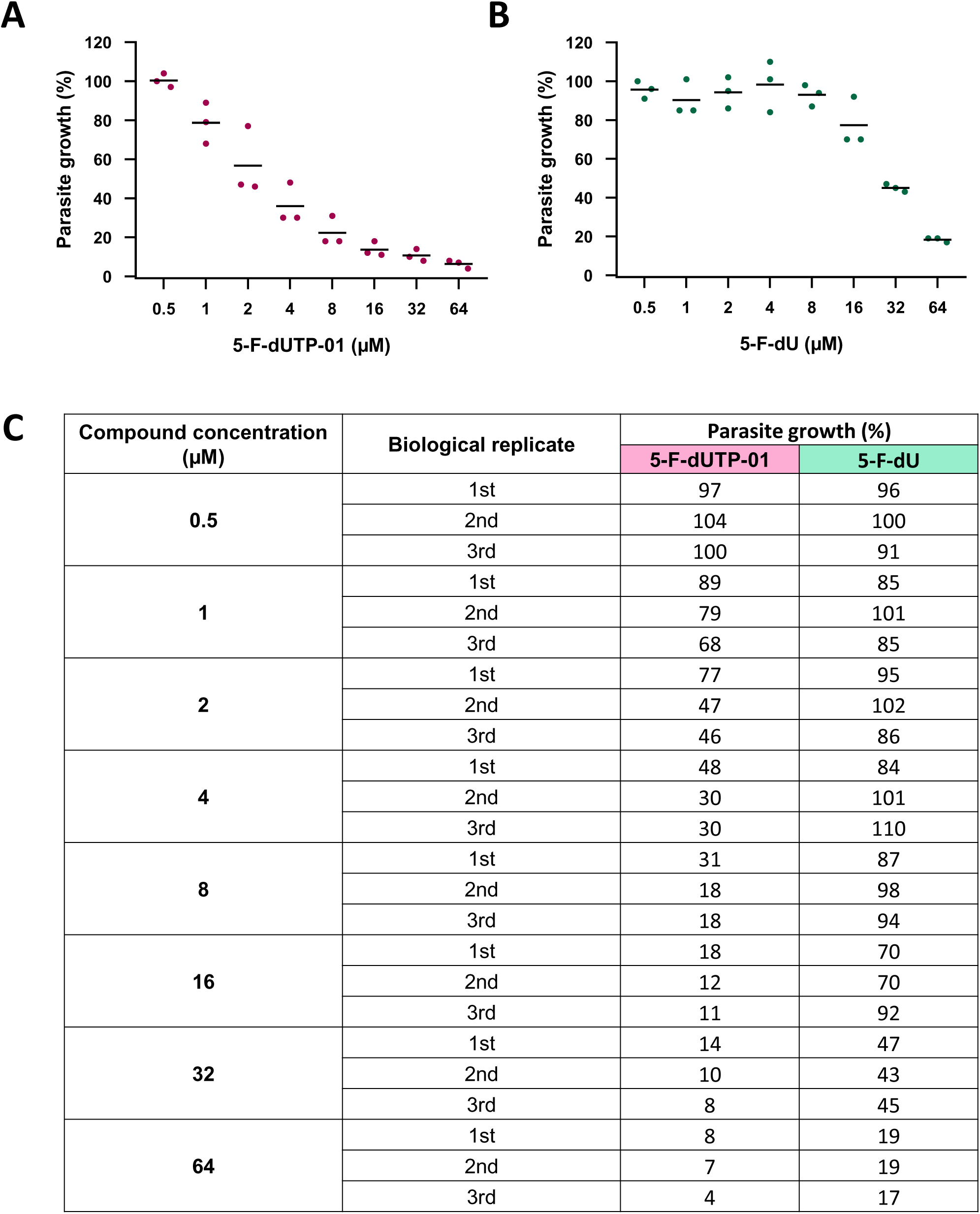
Impact of 5-F-dUTP-01 in comparison to 5-F-dU on the intraerythrocytic proliferation of *P. falciparum*. Corresponding to Figure 1B. **A.** Parasite growth in response to the indicated concentrations of 5-F-dUTP-01 was assessed by total DNA content measurement using the SYBR Green I fluorescence-based proliferation assay. The values obtained from each biological replicate were plotted. **B.** *P. falciparum* parasites were treated with 5-F-dU and their proliferation was evaluated as described in (**A.**). **C.** Quantification of parasite growth (percentage) from three biological replicates for both 5-F-dUTP-01 and 5-F-dU.

**Supplemental Figure 2:**
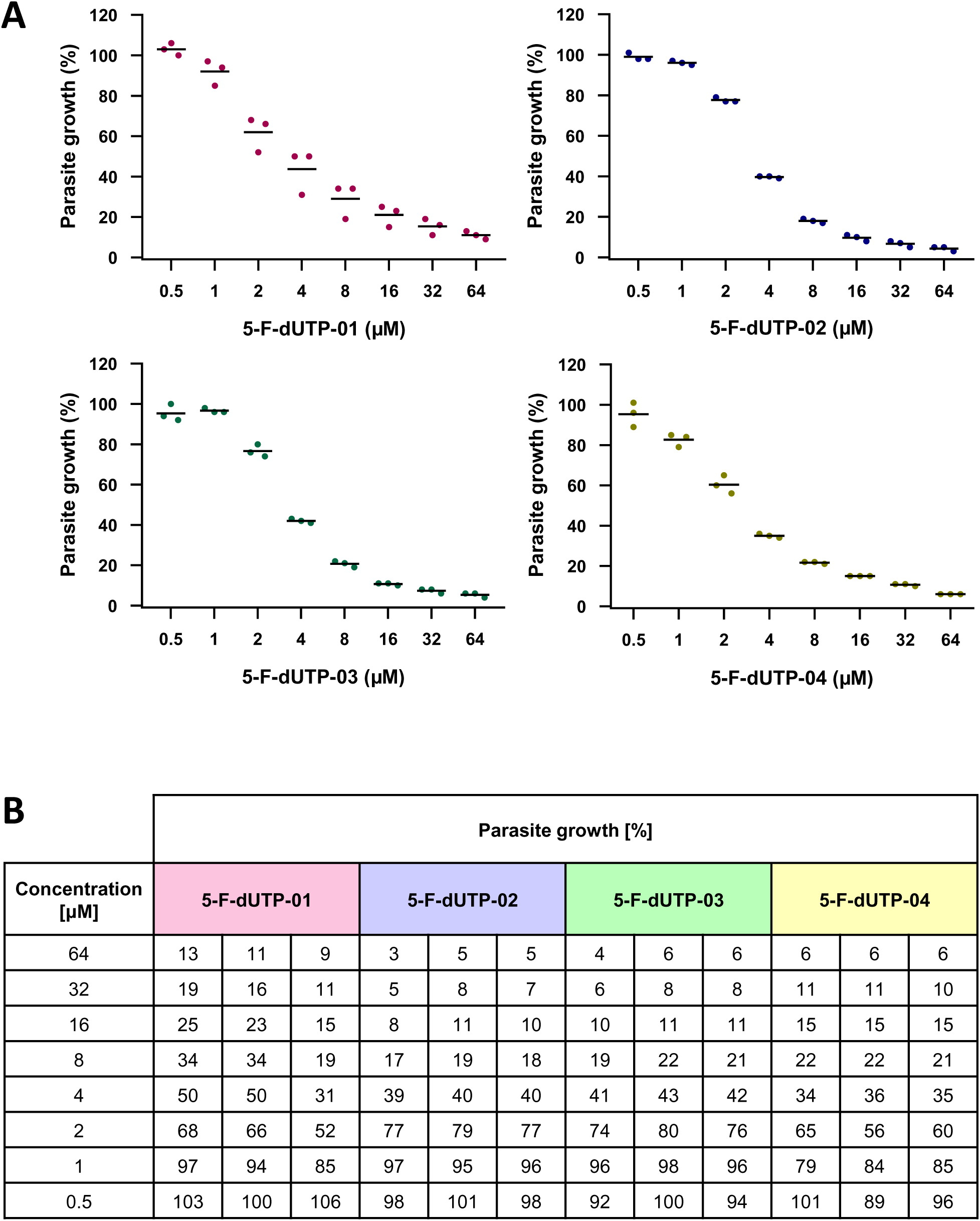
Parasite growth of *P. falciparum* 3D7 in response to cell-permeable FdUTP conjugates. Corresponding to Figure 1C. **A.** Parasite growth in response to cpFdUTPs was assessed by total DNA content measurement (SYBR Green I fluorescence-based proliferation assay). The values obtained from each biological replicate were plotted. **B.** Parasite growth in percentages, as compared to non-treated *P. falciparum*-infected RBCs. Each column represents a biological replicate. Based on this, the IC_50_ values were calculated by nonlinear regression (curve fit) using GraphPad Prism.

**Supplemental Figure 3:**
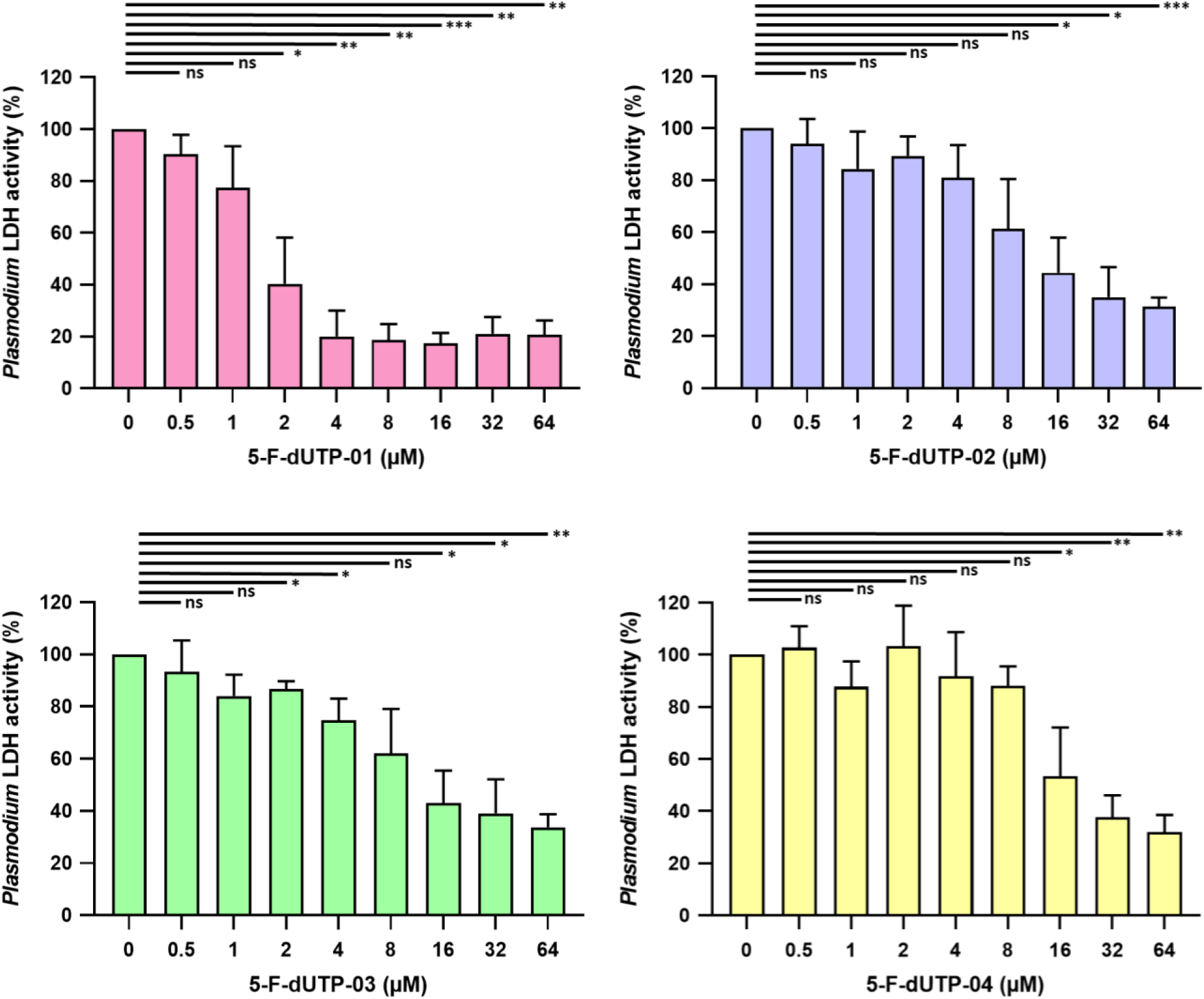
Impaired proliferation of *P. falciparum* in the presence of the FdUTP conjugates, assessed by *P. falciparum* lactate dehydrogenase activity. Corresponding to Figure 1C. **A.** *P. falciparum* was cultured in RBCs and then quantified through a colorimetric assay that specifically determines *P. falciparum* lactate dehydrogenase (*P. falciparum* LDH) but not mammalian LDH. The fraction of *P. falciparum* LDH activity observed in the presence of different cell-permeable FdUTP conjugates at different concentrations was determined and plotted. Statistical significance was assessed by t test with Welch’s correction, where a p-value <0.0001 was considered significant (*). ns = non-significant

**Supplemental Figure 4:**
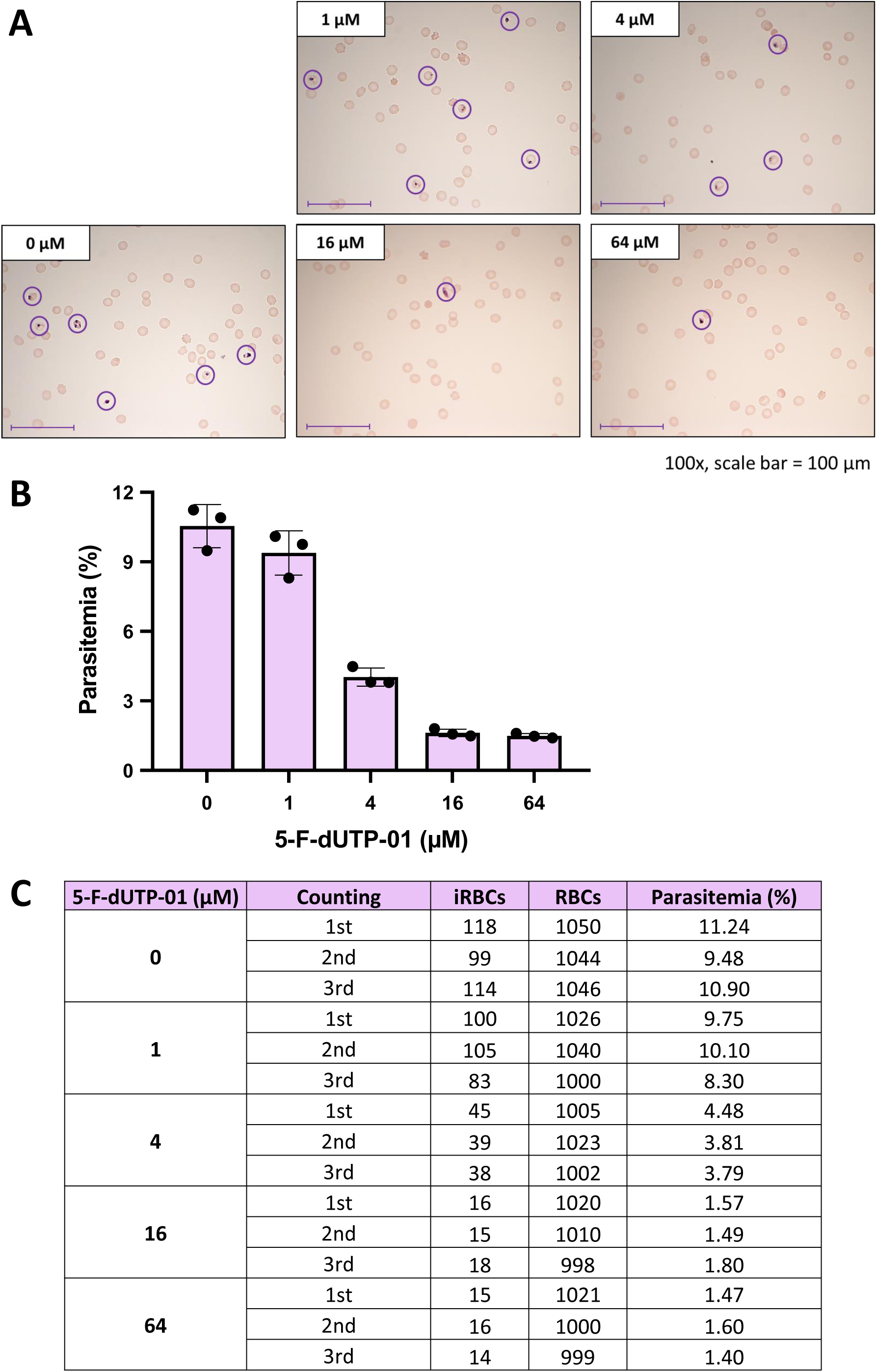
Impaired proliferation of *P. falciparum* in the presence of cpFdUTP (Giemsa staining). Corresponding to Figure 1B. Upon proliferation of *P. falciparum* for 72 hours in the presence of 5-F-dUTP-01 at the indicated concentrations, parasitemia was quantified by Giemsa staining. **A.** Representative images of Giemsa-stained *P. falciparum-*infected RBCs (circled) at varying concentrations of 5-F-dUTP-01. **B.** Quantification of the percentage of RBCs containing *P. falciparum* parasites (numerical replicates). **C.** Quantification of parasite growth from three rounds of counting.

**Supplemental Figure 5:**
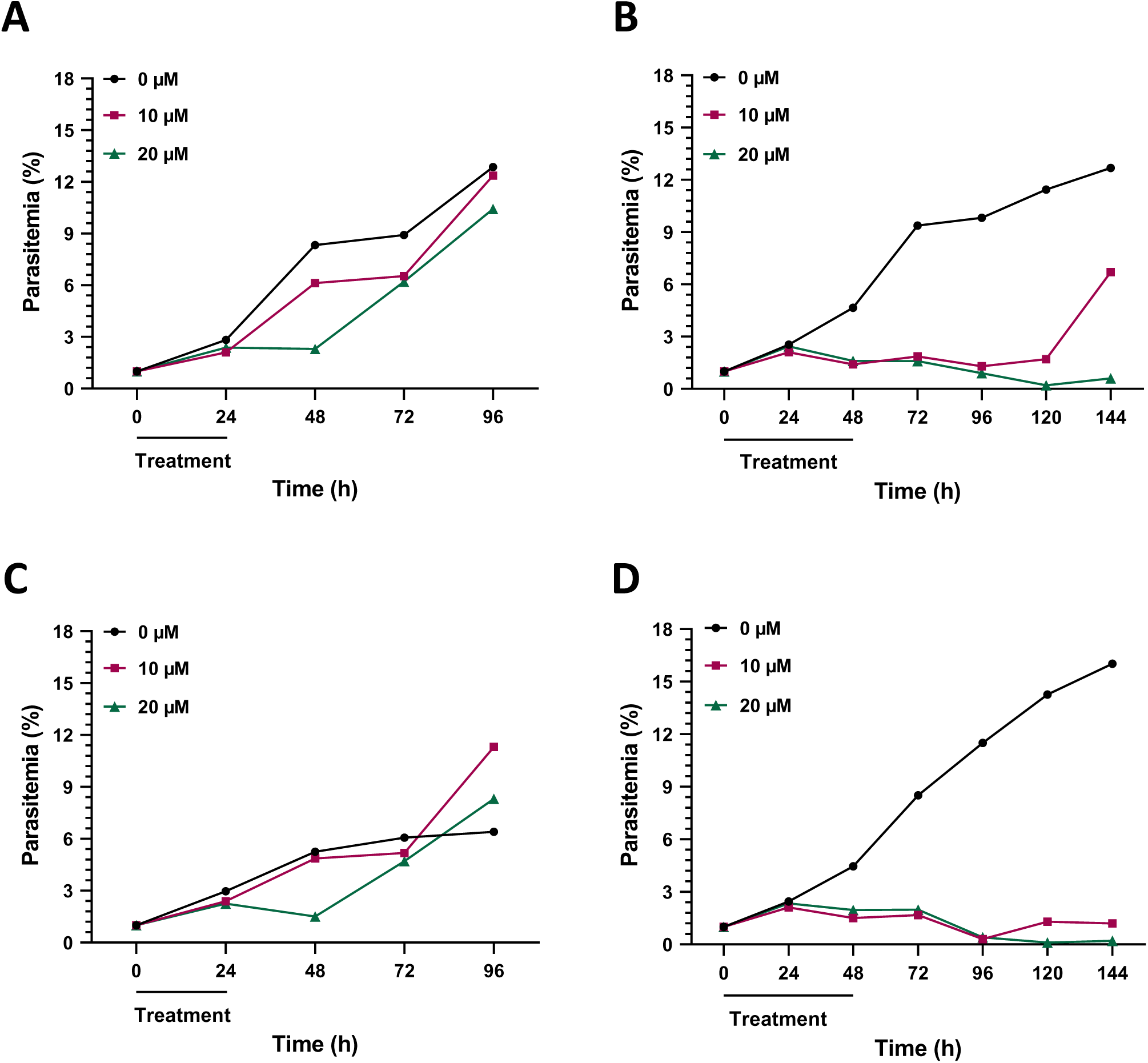
cpFdUTP inhibits *P. falciparum* growth in a sustainable fashion. Corresponding to Figure 2. **A.** *P. falciparum* 3D7 parasites were incubated with cpFdUTP at the indicated concentrations for 24 **(A, C)** or 48 (**B, D**) hours. After drug washout, the cultures were incubated further and parasitemia was determined by Giemsa staining. After 24 hours of incubation with the compound, parasites recovered quickly. Treatment with cpFdUTP for 48 hours inhibited the growth of *P. falciparum* in a sustainable fashion.

**Supplemental Figure 6:**
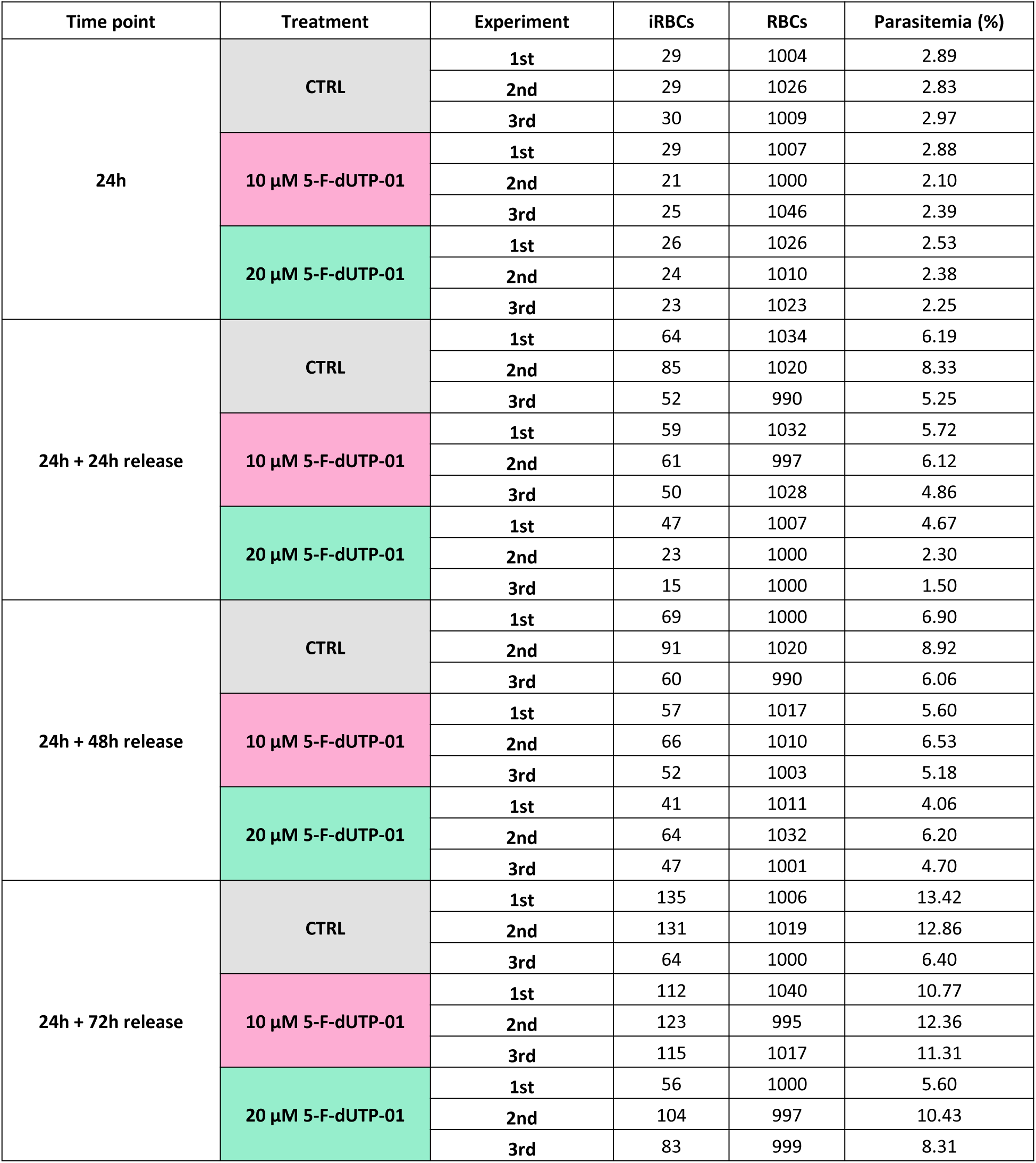
Raw data from the individual rounds of parasite counting in response to cpFdUTP after a 24-hour treatment. Corresponding to Figure 2A.

**Supplemental Figure 7:**
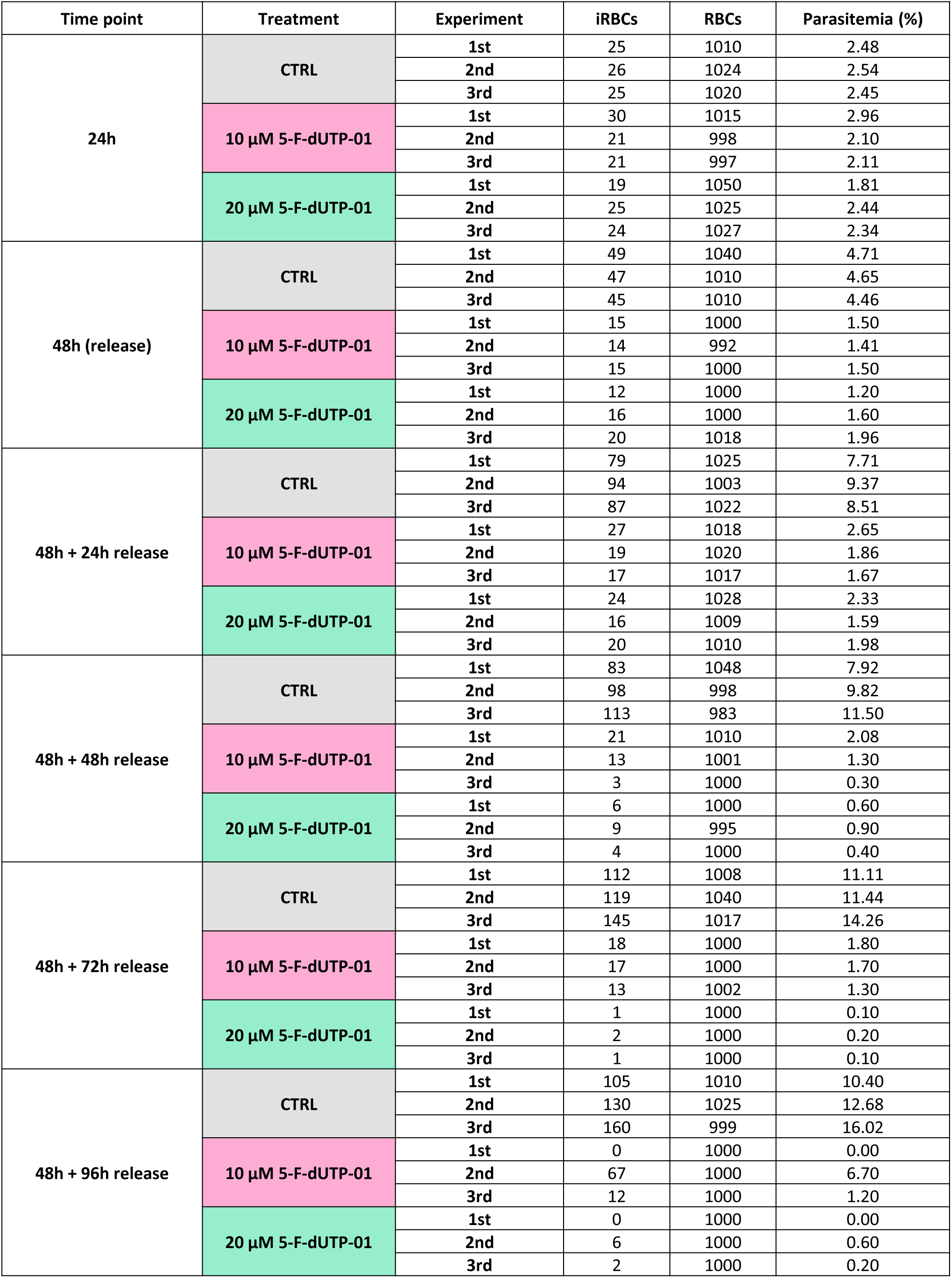
Raw data from the individual rounds of parasite counting in response to cpFdUTP after a 48-hour treatment. Corresponding to Figure 2B.

**Supplemental Figure 8.**
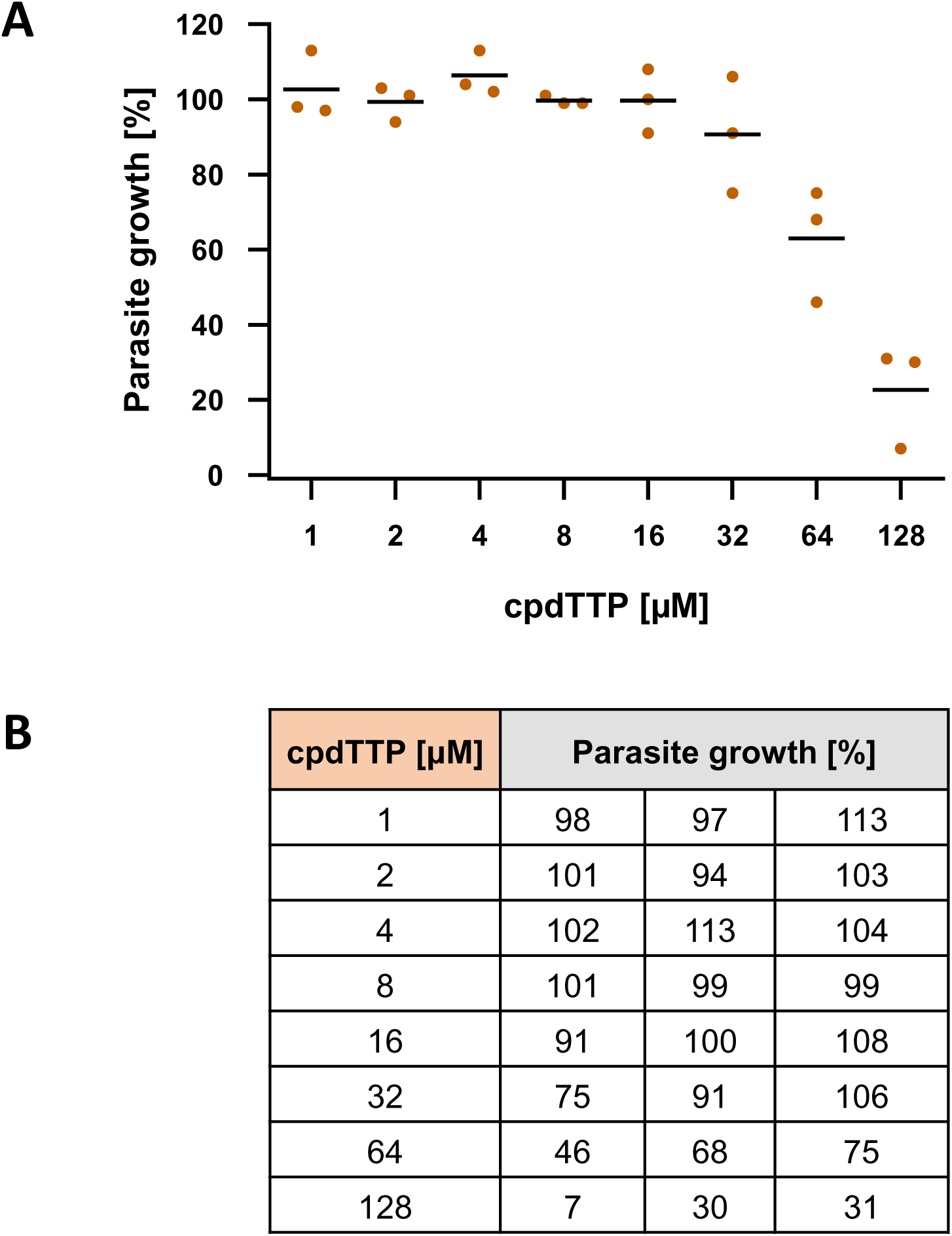
Little impact of cpdTTP on the proliferation of *P. falciparum*. Corresponds to Figure 3. *P. falciparum* was cultured as in Figure 1B while incubating the parasites with cell-permeable deoxythymidine triphosphate (cpdTTP) at increasing concentrations. Parasite proliferation was assessed by SYBR-Green staining of total parasite DNA. **A.** The values obtained from each biological replicate were plotted. **B.** Quantification of parasite growth. Each column represents a biological replicate.

**Supplemental Figure 9:**
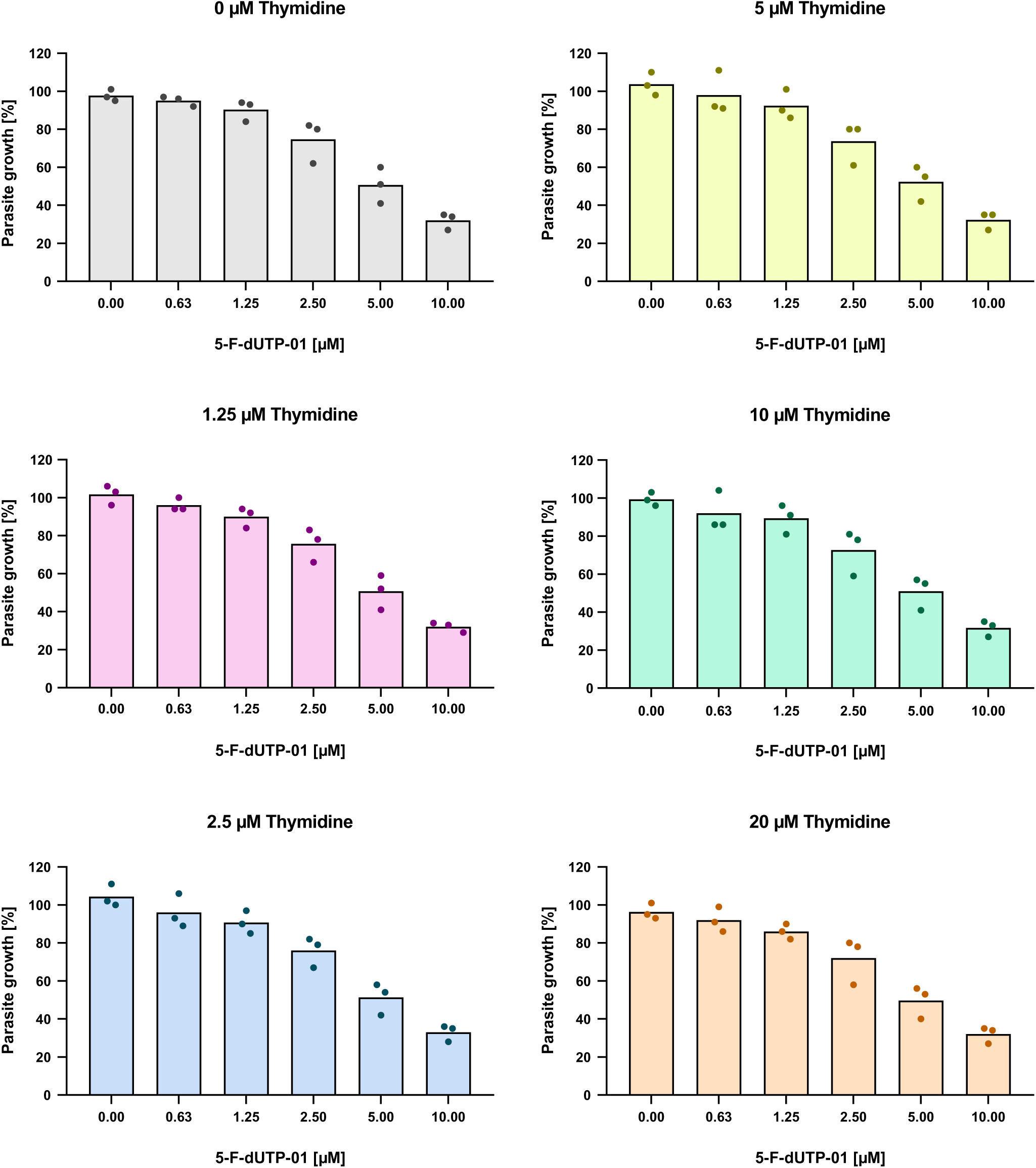
No ability of thymidine to rescue *P. falciparum* from cpFdUTP. Corresponding to Figure 4A. Asynchronously growing *P. falciparum* parasites were treated with cpFdUTP and thymidine at the indicated concentrations. After 72 hours, parasite growth was assessed by measurement of the total DNA content (SYBR Green staining). The values obtained from each biological replicate were plotted individually for each thymidine concentration.

**Supplemental Figure 10:**
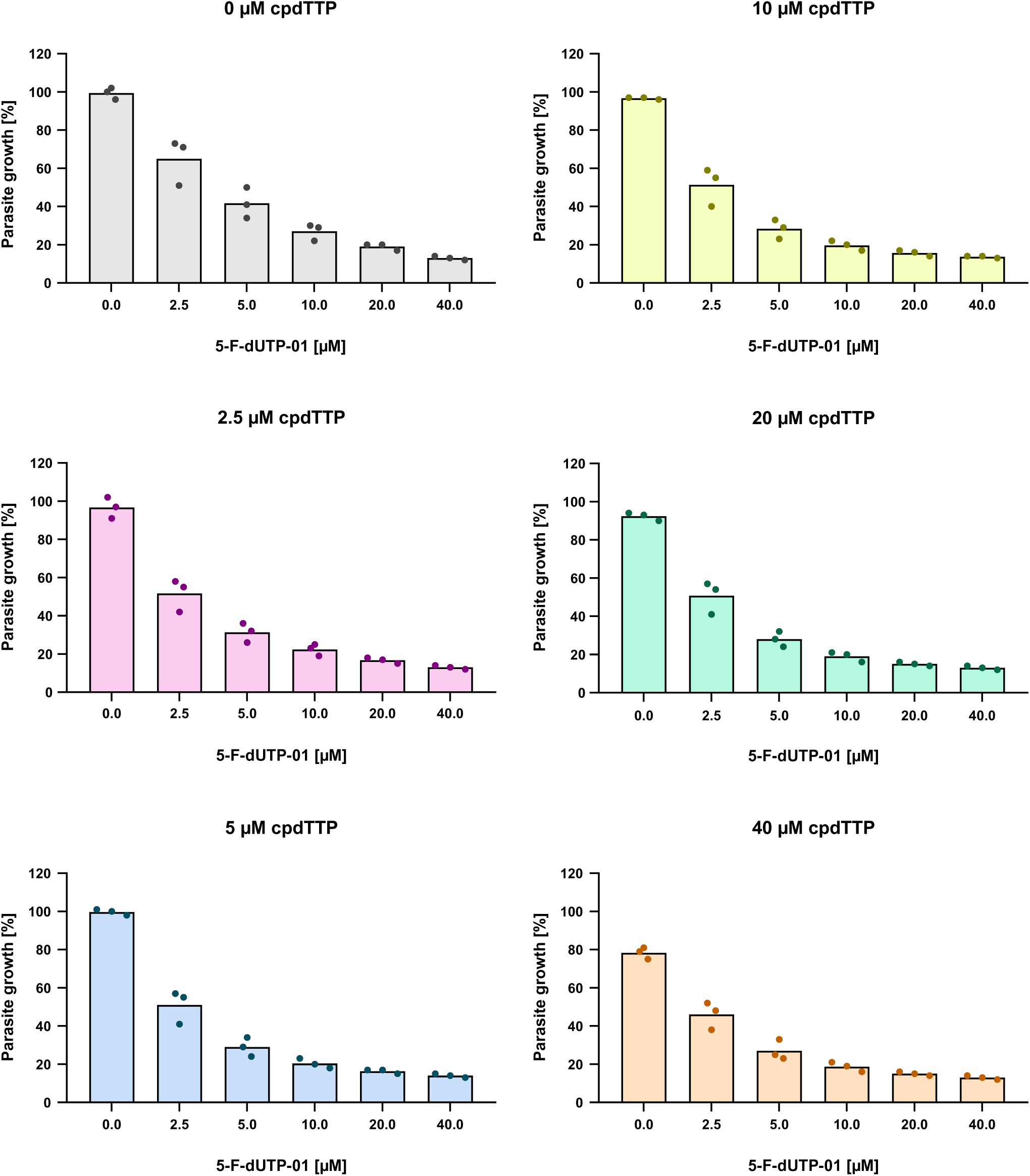
Failure of cpdTTP to rescue *P. falciparum* from cpFdUTP. Corresponding to Figure 4B. *P. falciparum* parasites were treated with 5-F-dUTP-01 in combination with cpdTTP. The values obtained from each biological replicate were plotted individually for each cpdTTP concentration.

***Supplemental Movie M1*: Prolonged nuclear localization of PCNA1 after exposure to cpFdUTP. Corresponding to Figure 6E**.

The experiment shown in Figure 6 was carried out with subsequent processing of images in a time-lapse fashion, resulting in a movie that visualizes strongly extended nuclear localization of PCNA1 when *P. falciparum* parasites were incubated with cpFdUTP (5-FdUTP-01).

